# Disruption of the HIV-1 Envelope allosteric network blocks CD4-induced rearrangements

**DOI:** 10.1101/827857

**Authors:** Rory Henderson, Maolin Lu, Ye Zhou, Zekun Mu, Robert Parks, Qifeng Han, Allen L. Hsu, Elizabeth Carter, Scott C. Blanchard, RJ Edwards, Kevin Wiehe, Kevin O. Saunders, Mario J. Borgnia, Alberto Bartesaghi, Walther Mothes, Barton F. Haynes, Priyamvada Acharya, S. Munir Alam

**Affiliations:** Department of Medicine, Duke University School of Medicine, Durham, NC 27710; Department of Surgery, Duke University School of Medicine, Durham, NC 27710; Department of Pathology, Duke University School of Medicine, Durham, NC 27710; Duke Human Vaccine Institute, Duke University School of Medicine, Durham, NC 27710; Genome Integrity and Structural Biology Laboratory, National Institute of Environmental Health Sciences, National Institutes of Health, Department of Health and Human Services, Research Triangle Park, NC 27709, USA; Department of Microbial Pathogenesis, Yale University School of Medicine, New Haven, CT 06536, USA; Department of Physiology and Biophysics, Weill Cornell Medicine, New York, NY 10021, USA; Department of Computer Science, Duke University, Durham, NC 27708; Department of Biochemistry, Duke University School of Medicine, Durham, NC 27710; Department of Electrical and Computer Engineering, Duke University, Durham, NC 27708; Department of Immunology, Duke University School of Medicine, Durham, NC 27710

**Author notes:** St. Jude Children’s Research Hospital, Department of Structural Biology, 262 Danny Thomas Place, Memphis, TN 38105-3678. Communicating authors (Rory Henderson, Priyamvada Acharya, S. Munir Alam).

## Abstract

The trimeric HIV-1 Envelope protein (Env) mediates viral-host cell fusion *via* a network of conformational transitions, with allosteric elements in each protomer orchestrating host receptor-induced exposure of the co-receptor binding site and fusion elements. To understand the molecular details of this allostery, we introduced Env mutations aimed to prevent CD4-induced rearrangements in the HIV-1 BG505 Env trimer. Binding analysis performed on the soluble ectodomain BG505 SOSIP Env trimers, cell-surface expressed BG505 full-length trimers and single-molecule Förster Resonance Energy Transfer (smFRET) performed on the full-length virion-bound Env confirmed that these mutations prevented CD4-induced transitions of the HIV-1 Env. Structural analysis by single-particle cryo-electron microscopy performed on the BG505 SOSIP mutant Env proteins revealed rearrangements in the gp120 topological layer contacts with gp41. Specifically, a conserved tryptophan at position 571 (W571) was displaced from its typical pocket at the interface of gp120 topological layers 1 and 2 by lysine 567, disrupting key gp120-gp41 contacts and rendering the Env insensitive to CD4 binding. Vector based analysis of closed Env SOSIP structures revealed the newly designed trimers exhibited a quaternary structure distinct from that typical of SOSIPs and residing near a cluster of Env trimers bound to vaccine-induced fusion peptide-directed antibodies (vFP Mabs). These results reveal the critical function of W571 as a conformational switch in Env allostery and receptor-mediated viral entry and provide insights on Env conformation that are relevant for vaccine design.

## Introduction

Host cell entry of HIV-1 is accomplished by the envelope glycoprotein (Env) spike. HIV-1 Env is a trimer of heterodimers comprised of gp120 and gp41 protomers, and exists in a metastable conformation capable of transitioning from a prefusion closed configuration to a fusion-competent open state upon triggering by CD4.^1,2^ The C-terminal gp41 domain contains a single transmembrane helix and the membrane fusion elements of the trimer.^3–5^ The gp120 segment binds the primary receptor CD4, triggering conformational changes leading to the binding of co-receptor CCR5/CXCR4 that causes global rearrangements in the trimer structure leading to viral and cell membrane fusion and gp120 shedding.^6–8^ While many studies have sought to understand the nature of the communication between the CD4 binding site, the coreceptor binding site and the fusogenic elements of the HIV-1 Env, a complete understanding of the allosteric mechanism and metastability in HIV-1 Env remains lacking.

The design of a soluble, stabilized ectodomain Env (SOSIP), containing an engineered gp120 to gp41 disulfide (SOS) and an HR1 helix breaking I to P mutation, has revealed structural details regarding broadly neutralizing antibody (bnAb) epitopes and as an immunogen has induced autologous neutralizing antibodies.^2,9^ The Env open state presents highly immunogenic, conserved fusion elements that typically induce poorly neutralizing antibody responses with limited heterologous breadth.^10^ Indeed, design efforts to improve the Env SOSIP by further stabilizing the closed Env conformation have resulted in multiple prefusion stabilized trimer designs capable of inducing improved autologous, difficult-to-neutralize tier 2 virus antibody responses.^10–21^ However, to date, no trimer design has successfully induced robust heterologous antibody responses.

Both the soluble and membrane-bound forms of the Env display an intrinsic ability to transition between multiple conformational states.^2,22–24^ In the pre-fusion, closed state, typical of SOSIP trimers, the gp120 domains surround a bundle of three gp41 helices, protecting conserved fusion elements of the trimer (Figure 1A). Interprotomer gp120 contacts exist at the trimer apex and form a cap that further encloses the gp41 three-helix bundle. This cap is composed of three sequence variable loop regions termed V1/V2 and V3 (Figure 1A).^3,9,25^ Global CD4-induced conformational changes result in dissociation of V1/V2 and V3 from the gp120 core toward a relatively disordered state as well as separation of gp120 from the gp41 three-helix bundle. This results in exposure of the three-helix bundle and fusion elements of the Env trimer.^26,27^ A layered architecture of the gp120 inner domain has been described^28^, with topological layers 1 and 2 contacting the gp41 subunit and shown, via mutagenesis coupled with cell-cell fusion and neutralization assays, to be important modulators of the CD4-bound conformation.^29^ Structures of the CD4-induced open state of the SOSIP Env, determined using cryo-electron microscopy (cryo-EM), showed that three gp120 tryptophan residues, W69, W112, and W427, form contacts from the CD4 binding site β20-β21 loop through to an intermediate helix, containing a portion of layer-2, to the distant gp120 layer-1 loop that is in contact with the gp41 three-helix bundle in the closed state (Supplementary Figure 1).^27^ In the open state structures, rearrangement of the layer-1 loop, which contains W69, further suggested that layer-1 plasticity plays a role in conformational transitions.^27^ Indeed, each loop has been implicated in the regulation of Env conformational transitions.^29,30^ Additional structural information from antibody-stabilized, potential closed-to-open state intermediates in SOSIP trimers suggests trimer opening occurs through ordered conformational transitions.^31^

**Figure 1.**
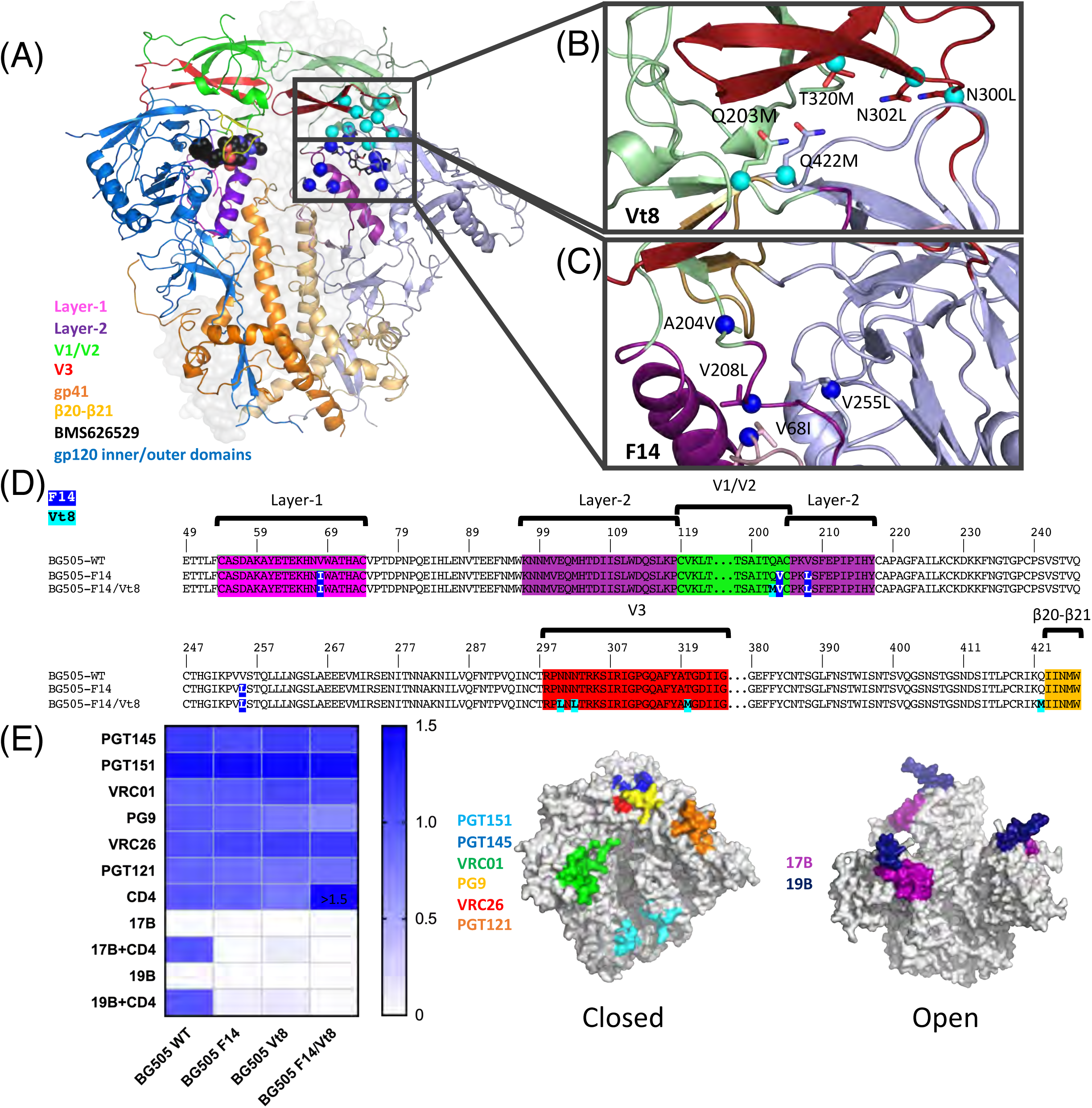
*Disrupting HIV-1 Env allostery to block CD4-induced conformational changes. **(A)*** HIV-1 Env SOSIP structure. Protomer to the left colored according to regions of allosteric control with BMS-626529 represented as black spheres. Protomer to the right in matte color scheme depicting the location of Vt-series mutations (cyan spheres) and F-series mutations (blue spheres). BMS-626529 depicted in stick representation. ***(B)*** Set of Vt8 residues. **(C)** Set of F14 mutations. Residue sidechains are represented as sticks with Cα atoms represented as cyan spheres. ***(D)*** Sequence alignment of BG505 WT, F14, and F14/Vt8 with F14 mutations highlighted in blue and Vt8 mutations highlighted in cyan. Layers 1 and 2, the V1/V2 region, and β20-β21 are highlighted using the same color scheme as in *(A)*. ***(E)*** Binding antigenicity of BG505 SOSIP.664 and mutants. *(left)* Heatmap of bnAb binding responses, CD4 binding, and CD4 triggering. Values for CD4 binding and CD4 triggering are normalized to BG505 SOSIP. *(middle)* Surface representation of a closed state SOSIP (PDB ID 5CEZ) and *(right)* an open state structure (PDB ID 5VN3). Colors identify antibody binding locations.

Despite these advances in our understanding of Env transitions, atomic level details of the allosteric mechanism by which CD4 induces transitions in the Env has remained elusive, thus limiting our ability to leverage such a mechanism for the development of vaccine immunogens. To investigate the allosteric mechanism of CD4-induced Env opening and to design SOSIP and membrane bound Env trimers that are resistant to CD4-induced structural changes, we aimed to interrupt the allosteric network responsible for CD4-induced triggering. The small molecule BMS-626529 is an attachment inhibitor that stabilizes the Env SOSIP in its prefusion closed state preventing CD4 binding and triggering.^32,33^ BMS-626529 neutralizes Env pseudoviuruses with nanomolar (nM) IC50, and has shown efficacy in blocking HIV-1 infection *in vitro* and *in vivo*.^32^ Further, BMS-626529 is known to stabilize membrane Env gp160 in its closed, functional state^33^, which is an important target in Env immunogen design efforts. Based upon the known allosteric control elements in the Env^27,28,30^ and atomic level details of the binding site of the BMS-626529 inhibitor^33^, we designed mutations that disrupted the Env allosteric network and rendered it unresponsive to CD4-induced structural rearrangements. Structure determination via single particle cryo-EM revealed key details regarding how allosteric elements downstream from the gp120-CD4 contact control transitions of the trimer between the Env closed and open states. These results provide a means by which to control the Env conformational ensemble and reveal new details required for understanding the conformational plasticity of the HIV-1 Env.

## Results

### Design of an Allosterically Decoupled, Conformationally Stabilized Env Construct

The small molecule HIV-1 entry inhibitor BMS-626529 has been shown to prevent sCD4-induced rearrangements in both soluble gp140 SOSIP trimers and native virion bound Env gp160.^32,33^ We reasoned that design of stabilized Env trimers based on an understanding of BMS-626529-mediated stabilization could lead to novel mutations that inhibit CD4 triggering for both the soluble SOSIP and membrane-bound Envs. A recent structure of a BG505 SOSIP in complex with BMS-626529 revealed that the compound resides in an induced pocket between the β20-β21 loop and the layer-2 α-1 helix, thus acting to separate the inner and outer domains (Figure 1A).^33^ The BMS-626529 compound appears to interrupt CD4 interaction by sequestering three key CD4 contact residues, N425, M426, and W427, thus impeding CD4 interaction and associated downstream rearrangements. Based upon the BMS-626529 contact region and neighboring residues in this structure, we selected clade A BG505 Env outer domain residues V255 and N377 and β20-β21 residues M426 and M434 for mutagenesis. (Figure 1A). In combination with these residues, additional sites were selected in layer-1 residues V68, H66, W69, and H72, layer-2 residue S115, and V1/V2 region residues A204 and V208 in order to prevent transitions in these conformationally plastic regions (Figure 1A). Together, this set of mutation sites, termed the F-series (Supplementary Table 1), included residues spanning the entire allosteric network region of gp120 from the CD4 binding site to the closed state site of gp120 contact with gp41 HR1. While the F-series mutations were designed to block CD4-triggering, we reasoned that V3 exposure may occur even in the absence of full triggering of the Env^34^ and examined residues in the V1/V2 to V3 contact region for mutagenesis to lock V3 in its prefusion, V1/V2-coupled state. We selected outer-domain residues E381 and Q422, V1/V2 region residues Q203, D180, and Y177, V3 residues R298, N300, N302 and T320, and residue Y435 for mutagenesis (Figure 1A). This set of mutations, termed the Vt-series (Supplementary Table 1), was introduced to prevent V3 exposure in a manner similar to previous stabilization strategies.^13,17^ Hydrophobic and/or space-filling mutations were made at each site for both the F-series and Vt-series sites. Beginning from a construct containing all sites in each series, smaller sets of mutations were prepared in order to examine the effect of particular mutations and to maximize the likelihood of identifying a suitable set of mutations for downstream processing (Supplementary Table 1). In order to examine their effect on the SOSIP trimer, each BG505 SOSIP mutant and the unmutated BG505 SOSIP (BG505 SOSIP) was transfected in HEK Freestyle293 (293F) cells followed by cell culture supernatant screening via biolayer interferometry (BLI) (Supplementary Figure 2A). The antibodies used in this screening included PGT145, 17B, 19B, and VRC01 in order to assess trimer quaternary conformation, CD4i epitope exposure, V3 exposure, and gp120 folding, respectively. Comparison of these results with the differences observed in the same assay for BG505 SOSIP in the presence and absence of BMS-626529 identified the ‘F’ series mutants F11, F14, and F15 as well as the ‘Vt’ series mutant Vt-8 as candidates for replicating the effects of the BMS-626529 compound. Each construct displayed a higher binding response to PGT145 and VRC01, no apparent 17B binding, and reduced 19B binding relative to the BG505 SOSIP (Supplementary Figure 2A; mutations from ^13^ included for comparison). The Vt8 mutations consisted of the V1/V2 mutation Q203M, V3 mutations N300L, N302L, and T320M, and the outer-domain mutation Q422M (Figure 1B and D). The F14 mutations consisted of the layer-1 mutation V68I, layer-2 mutations A204V and V208L, and the outer-domain mutation V255L (Figure 1C and D). The F15 mutant included the F14 mutations in addition to a gp120 outer-domain mutation N377L with F11 including mutations S115V, H72P, and H66S in addition to the F15 mutations. We selected the F14 construct for further characterization as it possessed the fewest number of mutations relative to F11 and F15. Since the F14 mutations were primarily in the layer 1 and 2 regions and were predicted to block the transition after CD4-induced destabilization of V1/V2 and V3, we combined F14 and Vt8 (F14/Vt8) in order to minimize V3 exposure (Figure 1D).

### Trimer Formation, Antigenicity, and Soluble CD4 (sCD4) Triggering of the Redesigned SOSIP Constructs

To evaluate the antigenicity of the designed Env mutants, we produced and purified the BG505 and mutant SOSIP Env variants by transient transfection in 293F cell culture followed by PGT145 affinity chromatography to select for well-folded trimers. The PGT145 purified material was further purified *via* size exclusion chromatography which resulted in a homogenous peak corresponding to the SOSIP trimer yielding a gp140 band when analyzed by non-reducing SDS-PAGE gel (Supplementary Figure 2B). We verified trimer formation for each construct by negative stain (NS) electron microscopy. The NS 2D-class averages of each construct confirmed that the mutants adopted a trimeric configuration similar to that of the BG505 SOSIP trimer (Supplementary Figure 2C).

We next examined the antigenicity of key bnAb epitope specificities for the redesigned SOSIPs *via* BLI using VRC01, PGT121, PG9, PGT145, PGT151, and VRC26 bnAbs, having CD4 binding site, glycan-V3, glycan-V1/V2, trimer apex, gp120/gp41 interface, and V1/V2 epitope specificities, respectively (Figure 1E). VRC01 binding indicated a ∼2-fold enhanced affinity for both Vt8 and F14/Vt8 relative to the BG505 SOSIP (Figure 1E, Supplementary Table 3, Supplementary Figure 3A). Fitting of the dose response curves for PGT121, PG9, PGT145, PGT151, and VRC26 indicated the mutations did not alter the affinity of the trimer for these important bnAb epitope specificities with nominal fold changes on the order of 1.0-1.3 (Figure 1E, Supplementary Table 3, Supplementary Figure 3A). These results indicated that the mutant designs presented a native, well-folded SOSIP trimer configuration and effectively presented multiple bnAb epitope specificities.

We next asked whether these mutations altered sCD4 binding. We determined the apparent affinity of each mutant construct and BG505 SOSIP for sCD4 *via* surface plasmon resonance (SPR). The affinities to CD4 determined for the F14 and Vt8 mutant SOSIPs matched that of BG505 SOSIP closely with K_D_s of 73.0 nM ± 26.2 nM, 83.9 nM ± 12.6 nM, and 67.9 nM ± 26.4 nM, respectively (Figure 1E, Supplementary Table 2, Supplementary Figure 3B). The BG505 F14/Vt8 SOSIP construct displayed a ∼4-fold enhanced CD4 affinity compared to BG505 SOSIP with a K_D_ of 15.6 nM ± 0.4 nM primarily as a result of an enhanced association rate (Figure 1E, Supplementary Table 2, Supplementary Figure 3B). As each construct bound CD4, we next asked whether sCD4 triggering was inhibited by the F14, Vt8, and F14/Vt8 mutations relative to the BG505 SOSIP. The CD4i antibody 17B and V3-targeting antibody 19B were used to monitor triggering of the coreceptor binding site and V3 exposed states, respectively. The results for the F14 construct indicated that sCD4 triggering of the open state is nearly eliminated based upon the lack of 17B response together with a ∼11-fold reduction in 19B response (Figure 1E, Supplementary Figure 3C). The Vt8 mutations similarly reduced 19B epitope exposure by ∼7-fold with triggering of the 17B epitope reduced by ∼9-fold (Figure 1E, Supplementary Figure 3C). Importantly, the combined F14/Vt8 construct eliminated CD4-induced exposure of both epitopes (Figure 1E, Supplementary Figure 3C). We also tested binding of the BG505 and mutant SOSIP Envs, in the presence and absence of sCD4, to a larger panel of V3 targeting MAbs, including 3074, 447-52D, and F39F. The results were consistent with those observed for 19B binding demonstrating triggering of BG505 SOSIP, reduced triggering of the F14 and Vt8 mutants, and little to no triggering of the F14/Vt8 mutant (Supplementary Figure 3F). To further validate these results, we measured 17B and PGT145 binding after incubation of BG505 and mutant SOSIP trimers with sCD4 for 30 minutes and 20 hours. The BG505 SOSIP displayed increased 17B binding after the 30-minute incubation with a further increase after 20 hours of incubation (Supplementary Figure 3D). The mutant Env SOSIPs were, however, little changed (Supplementary Figure 3D). Though the 30-minute incubation of BG505 with sCD4 had no effect on PGT145 binding, a marked reduction in binding was observed after 20 hours (Supplementary Figure 3E). This reduction was not observed in the mutant SOSIP Envs (Supplementary Figure 3E). Together, these results indicated that, while CD4 binding is retained, the mutant SOSIP Envs are not triggerable.

### Thermal Stability of F14 and F14/Vt8 SOSIPs

To assess the thermal stability of the F14, Vt8 and F14/Vt8 SOSIP trimers, and to compare them with BG505 SOSIP, we determined thermal denaturation maxima (T_max_) using differential scanning calorimetry (DSC), which showed that that the F14 mutations did not alter trimer thermal stability with T_max_ values of 66.3 °C ± 0.02 and 66.3 °C ± 0.06 for the BG505 and F14 constructs, respectively (Figure 2). The Vt8 and F14/Vt8 constructs displayed a 2.5 °C ± 0.02 and 1.8 °C ± 0.05 increase in T_max_, respectively, indicating the Vt8 mutations slightly improved thermal stability of the SOSIP trimer (Figure 2).

**Figure 2.**
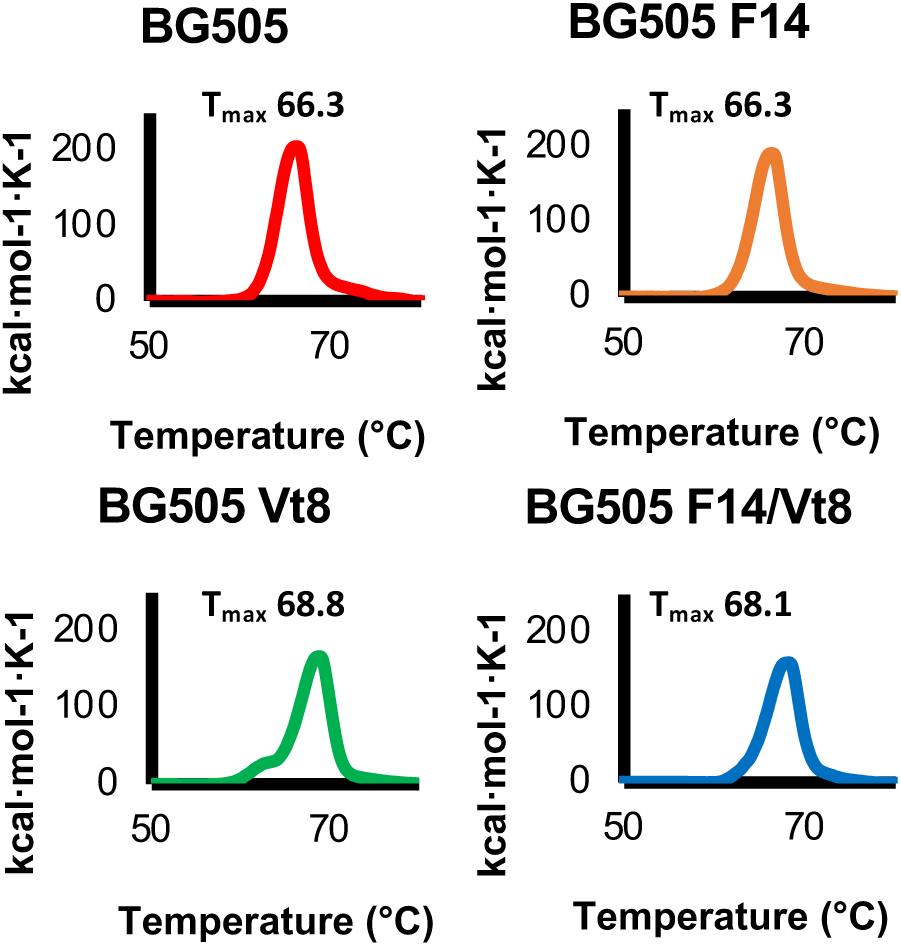
*Physical stability of BG505 SOSIP.664 and mutants.* Differential scanning calorimetry data for BG505 parent, F14, Vt8, and F14/Vt8 SOSIPs. Data presented are representative of three independent measures.

### Cryo-EM structures of the BG505 F14 and F14/Vt8 SOSIPs

To understand the structural basis for the observed lack of CD4-induced conformational rearrangements, we determined structures of the F14 SOSIP trimer in complex with VRC01 (Figure 3A) and the F14/Vt8 SOSIP trimer in complex with VRC03 and 10-1074 (Figure 3B) via single particle cryo-electron microscopy (cryo-EM). Map reconstruction was initially carried out in cryoSPARC^35^ followed by further refinement outside of cryoSPARC as described in the methods section (Supplementary Figure 4, Supplementary Table 4). A total of 77,632 and 84,378 particles yielded final map resolutions of 3.0 Å and 2.9 Å, respectively (Supplementary Figure 4). Fitting of atomic coordinates into each map revealed similar overall structures for both BG505 F14 and F14/Vt8 SOSIPs with a root mean square deviation of 0.70 Å for gp140 alignment (Figure 3C). The F14 mutations predominantly reside in V1/V2 near the apical region of layer-2 and in layer-1 near the gp120 contact with gp41 HR1 (Figure 1A-C, 3D-G). Clear densities for the F14 and Vt8 mutations were observed in both structures revealing minimal change in their positions relative to their typical SOSIP positions (Figure 3D and F, Supplementary Figure 5). However, map densities in the C-terminal portion of HR-1 in both F14 and F14/Vt8 trimers displayed a helical extension of the buried three helix bundle toward the trimer apex, a feature that resembles the extension observed in open and partially open SOSIP structures (Figure 3H, I, and J).^27,31^ As a result of this gp41 restructuring, residue K567 displaced W571 from the layer-1/layer-2 pocket that was formed by residues F43, C54, W69, T71, A73, C74, D107, L111, and T217 (Figure 4A and D). In the F14/Vt8 structure, this restructuring was associated with a rearrangement in layer-1 that reoriented residues H66 and H72 resulting in the formation of a potentially water mediated histidine triad configuration with gp41 HR1 H565 (Figure 4B, C, D). Interestingly, the F14 map displayed layer-1 loop densities suggestive of multistate behavior. Specifically, density corresponding to H66 and H72 in a configuration similar to that of the F14/Vt8 structure appeared alongside additional distinct densities, which may correspond to differing H72 sidechain configurations (Supplementary Figure 5). This suggested that the addition of the Vt8 mutations in the F14/Vt8 construct further stabilized the topological layer region facilitating the observed reduction in sCD4 triggering. Together, these results showed that structural changes in the region of gp120 contact with the C-terminal portion of gp41 HR1 caused by the F14 mutations effectively decoupled key allosteric control elements in the HIV-1 Env SOSIP trimer.

**Figure 3.**
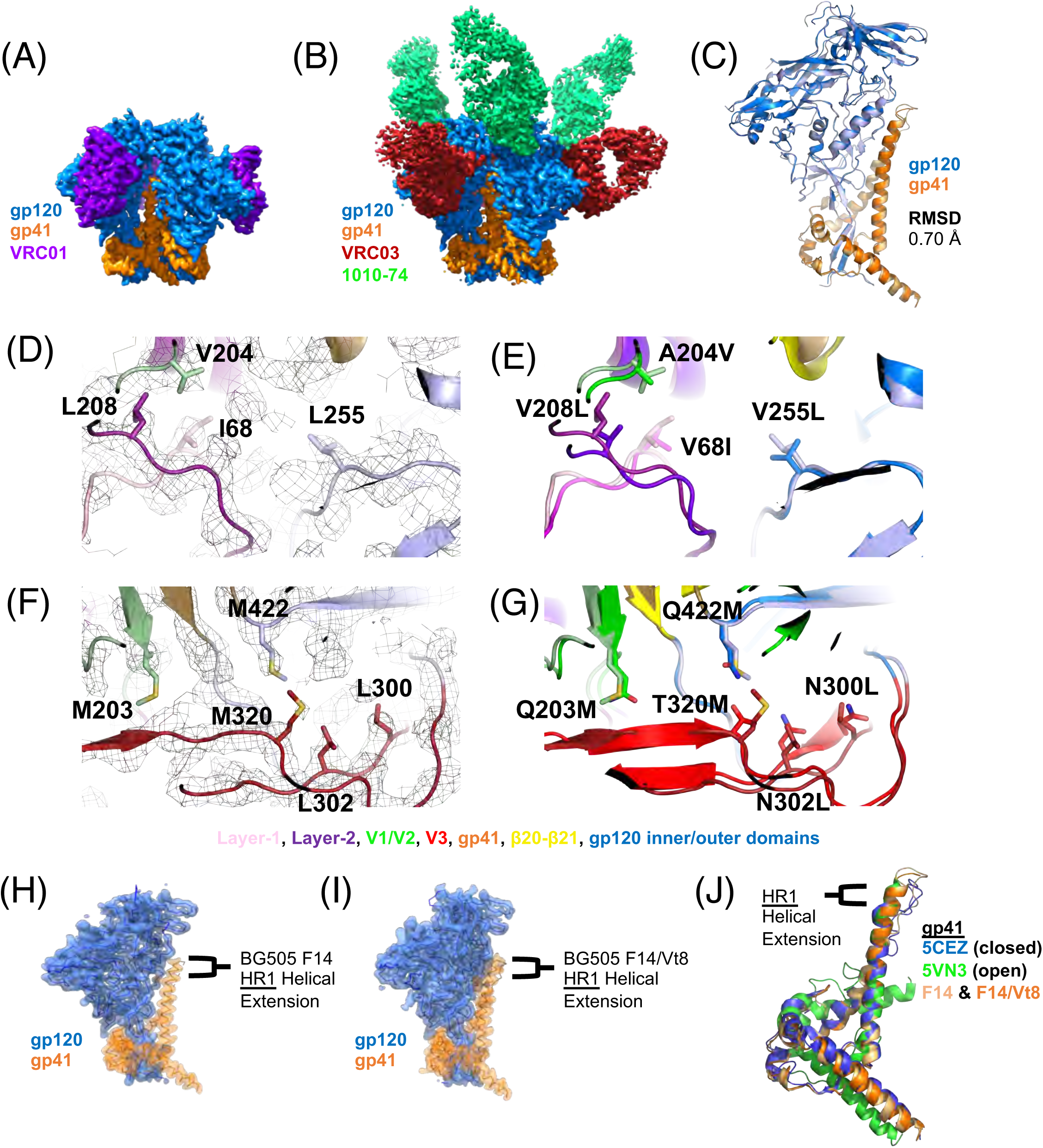
*Structural details of BG505 F14 SOSIP and BG505 F14/Vt8 SOSIP. **(A)*** Refined cryo-EM map of BG505 F14 SOSIP (gp120 in blue and gp41 in orange) bound to VRC01 (purple). ***(B)*** Refined cryo-EM map of BG505 F14/Vt8 SOSIP (gp120 in blue and gp41 in orange) bound to VRC03 (red) and 10-1074 (green). ***(C)*** Alignment of BG505 F14 SOSIP gp140 and BG505 F14/Vt8 SOSIP gp140. RMSD is for the alignment of gp120 and gp41 together. **(D)** Cryo-EM map of the BG505 F14/Vt8 construct with fitted coordinates depicting the F14 mutation sites. **(E)** Alignment of the BG505 F14/Vt8 SOSIP gp120 with the BG505 WT SOSIP gp120 showing the relative positions of the F14 mutations. **(F)** Cryo-EM map of the BG505 F14/Vt8 construct with fitted coordinates depicting the Vt8 mutations. **(G)** Alignment of the BG505 F14/Vt8 SOSIP gp120 with the BG505 WT SOSIP gp120 showing the relative positions of the Vt8 mutation sites. ***(H & I)*** Transparent cryo-EM maps with ribbon representations of refined coordinates for BG505 F14 and F14/Vt8 SOSIPs (gp120 in blue and gp41 in orange) identifying gp41 HR1 helical extension. ***(J)*** Alignment of gp41 three-helix bundle helix (residues 571-596) of BG505 F14 and F14/Vt8 SOSIPs (light orange and orange, respectively), an open state BG505 SOSIP (PDB ID 5VN3; green), and a closed state BG505 SOSIP (PDB ID 5CEZ; blue) identifying the extension of the gp41 HR1.

**Figure 4.**
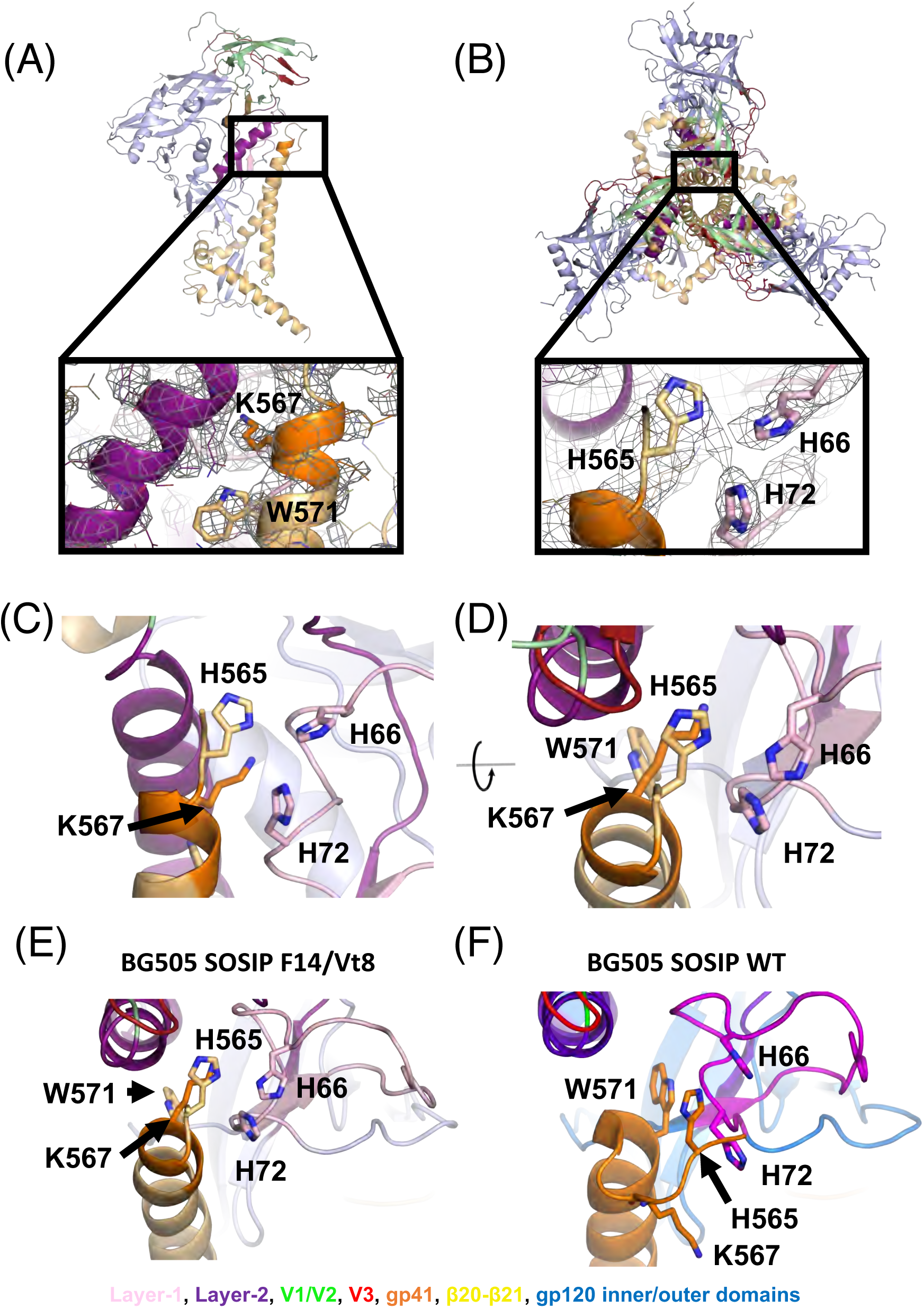
*Atomic level structural details of BG505 F14/Vt8 SOSIP. **(A)** (top)* Side view of VRC01 bound BG505 F14 SOSIP, with the rectangle highlighting the location of interior conformational change. (bottom) Zoomed-in view showing the gp41 region around residues K567 and W571 (shown as sticks), with the cryo-EM map shown as a mesh, and the underlying fitted coordinates in cartoon representation. ***(B)*** *(top)* VRC01-bound BG505 F14 SOSIP apex-facing trimer orientation, with the rectangle highlighting the location of interior conformational change. (bottom) Zoomed-in view showing the apparent histidine triad (shown as sticks) with the cryo-EM map shown as a mesh, and the underlying fitted coordinates in cartoon representation. ***(C)*** Side view of histidine triad relative to gp120 and residue K567. The viewing angles in Panels (B) and (C) are identical. ***(D)*** Top down view of histidine triad relative to gp120. ***(E & F)*** Comparison between the BG505 F14/Vt8 and WT SOSIP layer-1/2 and gp41 HR1 conformations.

### Comparison of F14 and F14/Vt8 to Previously Determined Env Trimer Structures

To understand the effect of the F14 and F14/Vt8 mutations on the overall structure of the SOSIP Env trimers, we examined regions of the structures distant from the F14 mutations. The individual domain coordinates of the mutant SOSIP trimer domains were found to be largely unperturbed indicating that the effects of the F14 and Vt8 mutations were localized. The gp120 domains within the Env Trimer are capable of rigid body movement relative to one another and to the gp41 three-helix bundle.^36^ We therefore devised a set of reference positions in gp120 and gp41 capable of describing structural rearrangements associated with rigid body movement in gp120 and gp41 (Figure 5). By comparing the closed^37^ and open state^27^ SOSIP trimer structures we identified two key points in the trimer about which the distance and angular disposition of the relevant domains could be described. Specifically, alignment of closed and open state (PDB IDs 5CEZ and 5VN3, respectively) gp41 residues G597-D664 that occur at the C-terminal end of the three-helix bundle helix demonstrated that the conformational transition from closed to open state yields a similar structure in this region with an RMSD of 0.721 Å (Figure 5A). This identified W596 as a hinge point about which the gp41 C-terminus rotates relative to the three-helix bundle helix. Comparing the position of the gp120 domains from this alignment revealed an additional trimer hinge-point between the gp120 N-terminal K46 and C-terminal K490 about which gp120 rotates (Figure 5B). We therefore devised a set of vectors connecting key points of the various trimer elements, including the W596 (A) and W571 (B) c-αs, a gp120 c-α centroid (C), a K46-K490 c-α centroid (D), and a V1/V2 c-α centroid (E), to enable comparison of the relative dispositions of the trimers in the closed, open and partially open (Open_p_) states (Figure 5C and Supplementary Figure 6A). Using these vectors, we performed a meta-analysis of Env structures deposited in the PDB. A total of 82 structures were analyzed including the newly determined F14 and F14/Vt8 structures (ref). We first examined the dihedral angle defined by the vectors connecting W571 c-α, W596 c-α, K46-K490 c-α centroid, and the gp120 centroid (BADC; Figure 5D). The dihedral angles calculated from the structures of the closed Env trimers clustered together with a mean of 10.5° (Figure 5D). Dihedral angles of the three partially open structures, PDB IDs 6CM3, 6EDU and 5THR, showed slightly shifted values of 0.7, −7.7°, and −16.0°, respectively, while the dihedral angles of the two open state structures displayed a marked shift at ∼ −87° each. The angles defined by W571 c-α, W596 c-α, and K46-K490 c-α centroid (BAD; named Angle 1) and by W596 c-α, K46-K490 c-α centroid, and gp120 centroid (ADC; named Angle 2) clustered in the closed state structures with means of 35.5° and 24.9°, respectively. Interestingly, the open state structures lie in the same closed state structure BAD angle cluster while the partially open state structures show a marked shift toward higher angles with a mean of 48.9°. When comparing the ADC angle, however, the open state structures, with a mean of 10.5°, differ from the closed state structures, while the partially open state structures have similar angles. The differences in the magnitude of the vector connecting the W571 c-α and the gp120 centroid were consistent with previous observations indicating that in both the partially open and open state structures the gp120 domains are shifted away from the gp41 core.^27,31^ These vector terms indicated that the transition from the closed state first involves a rotation about the W596 anchor away from the gp41 three-helix bundle followed by rotation about the vector connecting the W596 c-α and K46-K490 c-α centroid with a concomitant rotation in the direction of the three-helix bundle. The ability to discern closed, partially open, and fully open state structures confirmed that these vectors effectively reported on the relative disposition of gp120 to the gp41 three-helix bundle. We therefore examined whether the distance and angle terms between vectors connecting the key points, in combination with the dihedral terms, further characterized structural similarity between various antibody bound trimers. The spread in the angle and distance distributions limited direct analysis. Therefore, we employed the principal components analysis method to reduce the dimensionality of the dataset to enable clustering of similar SOSIP Env structures. Due to the increased number of terms and the possibility of fit uncertainty influencing the results, for this analysis we only used structures with reported overall resolution better than 4.5 Å. Additionally, the partially open and open state structures were excluded since they could dominate the PCs in this analysis. This resulted in the inclusion of a total of 59 structures. Clustering of the PCA results indicated that the F14 and F14/Vt8 structures indeed differed from the majority of the available SOSIP structures, residing in distinct clusters closer to a cluster of fusion peptide-directed antibody-bound SOSIP structures including those induced in mice and Rhesus macaques (Figure 5E, Supplementary Table 5).^38^ This finding indicated that the changes observed in the layer1/2 contact region with gp41 indeed altered the overall orientation of gp120 relative to gp41, and showed that these changes were structurally similar to those induced by fusion peptide-directed antibodies. We next examined the structure of the vFP antibodies in the region of the F14 conformational changes. All four mouse vFP-bound, stabilized BG505 SOSIP trimers displayed density consistent with the helical extension and W571 rearrangement suggesting the rearrangement in the F14 and F14/Vt8 structures represented a functionally accessible state of the Env and that vFP antibodies may neutralize via inhibition of HIV-1 Env triggering (Figure 5F, Supplementary Figure 6). The F14 cluster contained one vFP bound structure and a BG505 RC1 SOSIP^39^, designed to enhance vaccine induced V3-glycan responses, in complex with 10-1074 and a CD4bs Fab. Interestingly, antibodies in the F14/Vt8 cluster included the RC1 SOSIP bound to vaccine induced V3-glycan targeting Fabs and the partially open state stabilizing 8ANC195 Fab suggesting the 8ANC195 Fab shifted the configuration relative to the 10-1074 bound structure. The remaining structure in the F14/Vt8 cluster was for a CH505 SOSIP Env bound to the CH235 UCA Fab.^40^ Examination of the CH505 Env density near the gp41 HR1 C-terminus indicated W571 displacement as in the F14 and F14/Vt8 structures. Density in the RC1 structures was not sufficiently clear to determine the extent to which this region may differ from the typical SOSIP structure. Together, these results demonstrated the F14 and F14/Vt8 mutations induced global rearrangements in the SOSIP that may represent an accessible state to the native Env.

**Figure 5.**
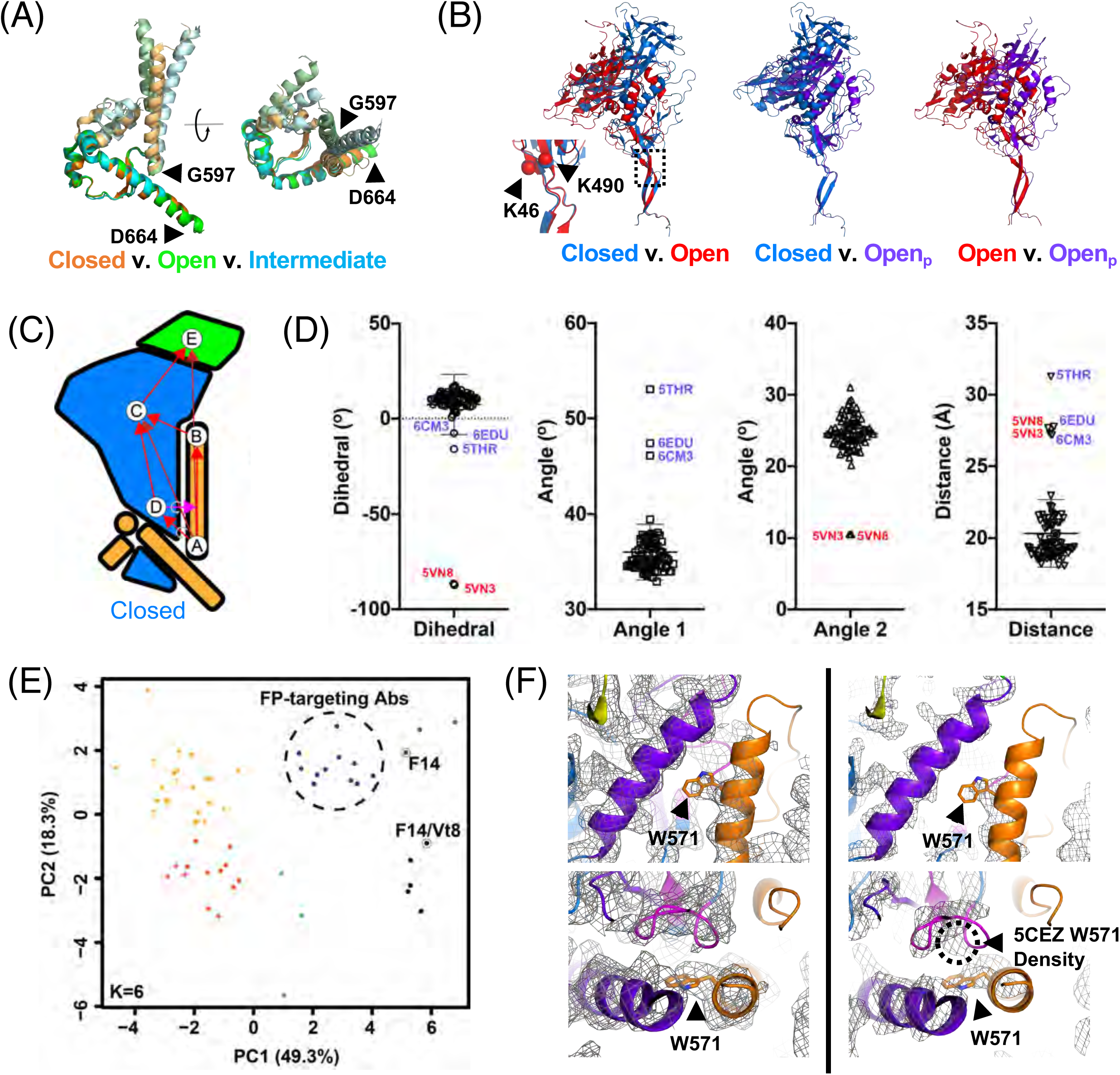
*Domain organization and antigenicity of the BG505 F14 and BG505 F14/Vt8 SOSIP **(A)*** Alignment of gp41 residues 597 to 664 in the closed (PDB ID 5CEZ), partially open (Open_p_; PDB IDs 6CM3 and 6EDU), and open (PDB IDs 5VN3 and 5VN8) SOSIP trimers. ***(B)*** Relative orientations of gp120 in the closed (PDB ID 5CEZ), partially open (PDB IDs 6CM3), and open (PDB IDs 5VN3) SOSIP trimers aligned on gp41 residues 597 to 664. *(inset, left)* Relative gp120 orientations from the gp41 597 to 664 residue alignment depicting the location of the K46/K490 hinge-point. Note: the color coding for the SOSIPs here differs from that in panel (*A*). ***(C)*** Cartoon representations of gp120 (light blue), gp120 V1/V2 region (green), and gp41 (yellow) in the closed state. Letters indicate location of centroids with arrows depicting vectors between centroids. **(D)** (left) BADC dihedral angles for closed, partially open, and open state structures. (middle left) BAD angles for closed, partially open, and open state structures. (middle right) ADC angles for closed, partially open, and open state structures. (right) BC vector magnitudes for closed, partially open, and open state structures. Open state structures (PDB IDs 5VN8 and 5VN3) are marked with red labels, and partially-open structures (PDB IDs 6CM3, 6Edu and 5THR) are shown with purple labels. ***(E)*** Principal components analysis of centroid vectors with principal component one and two scores plotted for each closed state SOSIP analyzed. Points are colored according to k-means clusters (K=6; associated PDB IDs: Supplemental Table 5). ***(F)*** *(left)* Alignment of the BG505 F14/Vt8 coordinates with the vFP7.04 bound BG505 DS SOSIP cryo-EM map (EMD-7621). *(right)* Alignment of the BG505 F14/Vt8 coordinates with the PDB ID 5CEZ map. W571 (sticks) aligns with density corresponding to 5CEZ V570 coordinates.

### Antigenicity, CD4 triggering, and Conformational Distribution of the Redesigned gp160 Trimers

To determine whether the effects of the F14/Vt8 mutations observed in the soluble SOSIP trimers translated to native, membrane-bound gp160 trimers, we used two measures: 1) we assessed the antigenicity and CD4-triggering of cell surface Env gp160s using 293F cells displaying full length trimers on their surface and 2) we performed smFRET experiments on BG505 Env on the virion surface.

In the assay using cell surface Env gp160s, BG505, a previously engineered BG505 stabilized design termed BG505 DS^25^, and BG505 F14/Vt8, BG505 F14, and BG505 Vt8 were first tested for binding to bnAbs N6, CH01, PGT125, and PGT145 targeting CD4bs, V1/V2, glycan-V3, and trimer apex epitopes, respectively. Binding to cell surface expressed Env trimer was assessed via flow cytometry to determine the percentage of positive cells and mean fluorescence intensities (MFI) for bnAb binding (Supplementary Figure 7). A number of factors may influence the observed binding including the presence of misfolded or degraded Env. We therefore primarily focus on the relative differences between constructs and conditions. The percentage of cells testing positive for binding to each bnAb were similar in each construct with the exception of BG505 DS binding to CH01, which displayed a marked reduction in cells positive for CH01 binding (Figure 6A, Supplementary Figure 8A, Supplementary Table 6). We next examined binding of non-bnAbs 19B and 17B, which target the open state V3-tip and bridging sheet epitopes, respectively. Binding to these open state preferring antibodies was tested in the presence and absence of sCD4 or potently neutralizing eCD4-Ig, a coreceptor-mimetic peptide fused eCD4-Ig^41^. In order to ensure comparable levels of Env surface expression between constructs, the near-pan neutralizing bnAb N6^42^ was used as a benchmark yielding comparable binding for all constructs tested with MFIs of 457.3 ± 137.9, 338.0 ± 72.6, 341.3 ± 43.2, 568.0 ± 152.6, and 378.3 ± 73.1 for BG505, BG505 DS, BG505 F14/Vt8, BG505 F14, and BG505 Vt8, respectively (Supplementary Figure 8B and C, Supplementary Table 7). Pairwise percentage positive cell comparison of each construct in each replicate experiment demonstrated a consistent reduction of intrinsic exposure of the 17B open state epitope in both the BG505 DS and BG505 F14/Vt8 stabilized gp160 trimers as compared to BG505 gp160 (Figure 6B, Supplementary Figure 8D and E, Supplementary Table 6). Intrinsic exposure of the V3 tip targeting 19B epitope was relatively similar between BG505 and BG505 DS gp160s while the BG505 F14/Vt8 gp160 displayed a consistent ∼2-fold reduction in 19B epitope exposure relative to BG505 gp160 (Figure 6B, Supplementary Figure 8D and E, Supplementary Table 6). On incubating the cells with either sCD4 or eCD4-Ig, distinct increases in both the 17B and 19B percentage of positive cells were observed for BG505 gp160, as expected (Figure 6B, Supplementary Figure 8D and E, Supplementary Table 6). Conversely, binding to these CD4-induced epitope-targeting antibodies was not observed for BG505 DS or F14/Vt8 stabilized gp160 trimers in the presence of either sCD4 or eCD4-Ig (Figure 6B, Supplementary Figure 8D and E, Supplementary Table 6). Triggering of the 17B epitope was not observed in BG505 F14 or BG505 Vt8 gp160s in the presence of sCD4, although 19B epitope exposure was observed in each (Supplementary Figure 8F and G, Supplementary Table 6). Additionally, both 17B and 19B epitope exposure was observed in the presence of eCD4-Ig (Supplementary Figure 8F and G, Supplementary Table 7). These results showed that the F14, Vt8, and F14/Vt8 mutations allowed presentation of key bnAb epitopes and that the combined F14/Vt8 mutations were necessary for the minimization of intrinsic non-bnAb epitope exposure and the prevention of CD4 induced rearrangement of the surface expressed gp160 Env.

**Figure 6.**
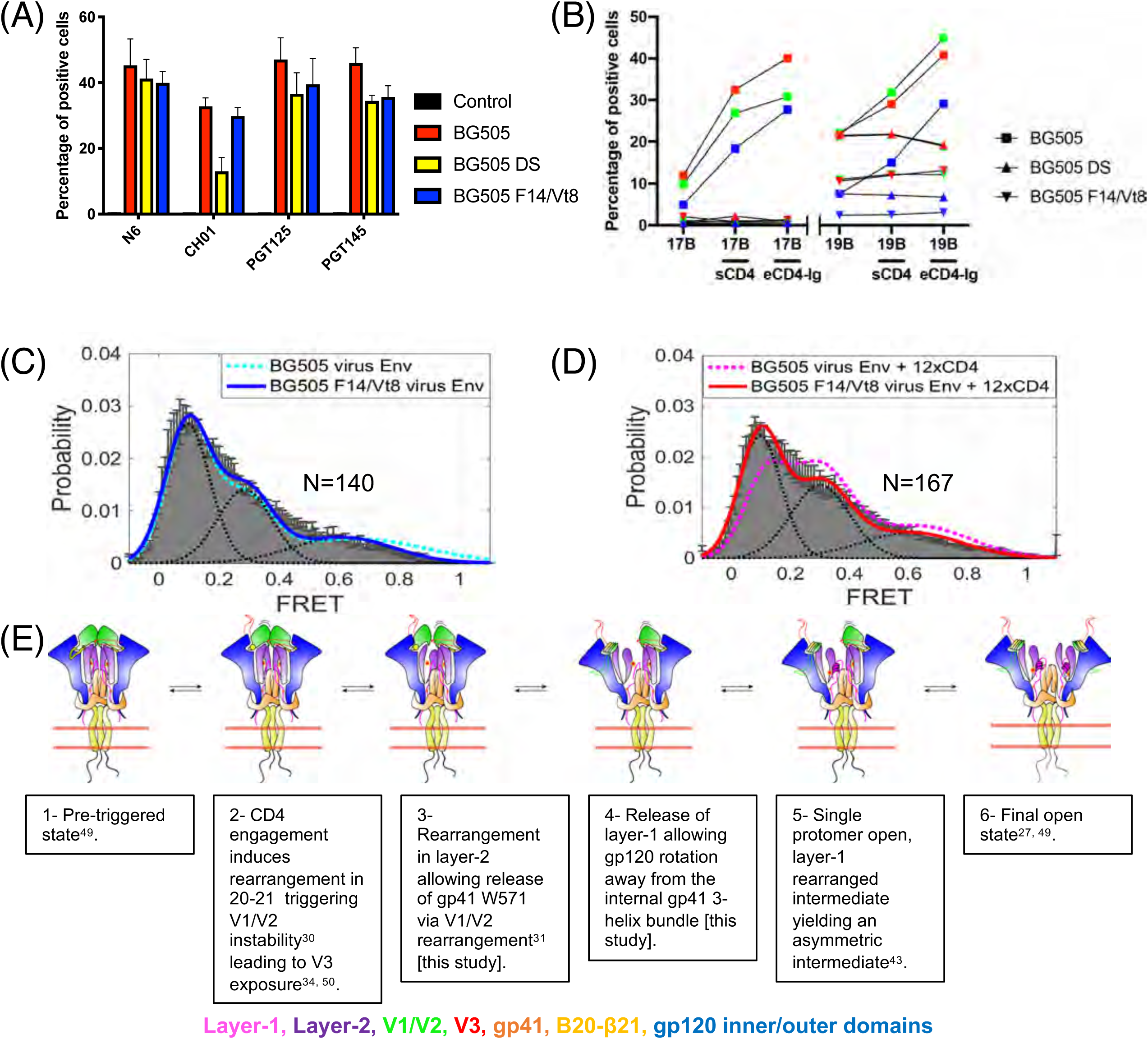
*F14/Vt8 mutations stabilize native, membrane bound Env. **(A)*** Percentage of cells positive for binding of bnAbs N6, CH01, PGT125, and PGT145 to 293F cell surface expressed BG505, BG505 DS, and BG505 F14/Vt8 trimers. ***(B)*** Chart of the percentage of positive cells from each independent triggering experiment colored according to the first (red), second (green) and third (blue) experiments with BG505 represented as squares, BG505 DS represented as upward oriented triangles, and BG505 F14/Vt8 represented as downward oriented triangles. ***(C)*** Single molecular FRET distribution for BG505 WT (dashed cyan line) and F14/Vt8 (solid blue line) virus Env. ***(D)*** Single molecular FRET distribution for BG505 WT (dashed pink line) and F14/Vt8 (solid red line) virus Env in the presence of dodecameric CD4. ***(E)*** Mechanism for transition between a closed state trimer to an open state trimer.

We also used the smFRET assay developed by Munro et. al^24^, to examine the conformational landscape and CD4-induced effects on the virion bound F14/Vt8 Env as compared to the BG505 Env. Specifically, we examined the FRET distributions of the BG505 and F14/Vt8 BG505 Envs to determine a) whether the mutations alter the unliganded FRET distribution relative to BG505 Env and b) whether the mutations prevent dodecameric CD4 (sCD4D1D2–Igαtp) induced rearrangements. smFRET studies previously revealed that native Env on virus predominantly resides in in a conformational state termed State 1 but has spontaneous access to two more conformational states, termed states 2 and 3. In response to CD4 binding, wild-type Env opens into State 3 through one necessary intermediate (State 2).^24,43^ Comparison of the unbound Envs indicated that both BG505 and BG505 F14/Vt8 unliganded Envs displayed similar distributions, with the predominant State 1 peak at low FRET, yielding State 1 occupancies of ∼46% and ∼49% for the BG505 and F14/Vt8 Envs, respectively (Figure 6C and D, Supplementary Table 8; example FRET traces shown in Supplementary Figure 9). Both trimers also displayed similar occupancies of State 2 and State 3. Addition of CD4 resulted in a shift in the distribution for the BG505 Env toward increased high and mid-FRET states corresponding to the asymmetric intermediate State 2 configuration and the open State 3 configuration, respectively (Figure 6C and D, Supplementary Table 7). Unlike the BG505 Env, however, the F14/Vt8 Env did not respond to addition of dodecameric CD4, displaying the same relative distribution of the three FRET states in the presence of CD4 as were seen in the absence of CD4, and consistent with the observed resistance to CD4-induced changes in the BG505 F14/Vt8 SOSIP Env (Figure 6C and D, Supplementary Table 7). Together, results for the cell and viral membrane associated gp160 Env F14/Vt8 mutant trimer are consistent with observations for these mutations in the soluble SOSIP trimer indicating the mutations designed to disable Env allostery effectively blocked CD4-induced rearrangements while maintaining efficient bnAb interaction.

## Discussion

The HIV-1 Env is an intricate conformational machine that propagates receptor-mediated structural changes from a neutralization-resistant closed conformation to a fusion-competent open conformation that exposes immunodominant epitopes for non-neutralizing antibodies. The closed conformation of the HIV-1 Env is of interest for vaccine design since it presents the epitopes for broadly neutralizing antibodies, and the expectation is that effective presentation of the native, closed conformation of the Env by vaccination is essential to elicit broadly neutralizing antibodies. Indeed, many studies have succeeded in stabilizing the closed conformation of the HIV-1 Env both in the soluble Env format, as well on the cell surface, with varying levels of success in eliciting autologous and heterologous neutralization.^10–21^

In this study we combined structural and mechanistic information to construct a mutant Env that is no longer responsive to triggering by the CD4 receptor. Two strategies were employed, each aimed at shifting the equilibrium of Env dynamics towards its closed state. The first strategy that led to the F14 series of mutations identified the conserved W571 as a conformational switch and effectively disabled a communication network that relied on the movement of topological layers 1 and 2 to transmit structural changes from the CD4 binding site. The second path aimed to prevent V3 exposure via mutation of buried hydrophilic to hydrophobic residues in order to prevent the infiltration of water into the space between V3 and V1/V2. This second strategy was similar to recent stabilization strategies also aimed at preventing V3 exposure that yielded similar results.^10,13,17^ Comparison of the F14 and Vt8 designs with previous stabilization strategies via the vector-based analysis (Figure 5D and E) revealed that most stabilized SOSIP structures, with the exception of those bound to fusion peptide-directed antibodies and the RC1 structures, resided in two dominant closed state clusters (Figure 5E; Supplementary Table 5). Interestingly, unlike previous stabilization strategies that either reduce or prevent CD4 binding^12,13,17,44^, the F14/Vt8 trimer retained interaction with CD4 while still preventing CD4-induced exposure of open state epitopes. Together, the findings presented here indicated close coupling of sCD4 induced internal rearrangements in gp120 and the N-terminal portion of the gp41 three-helix bundle.

While the F14 and Vt8 designs were made and tested in the context of the soluble SOSIP Env, the observed effects translated to the Env on the native virion surface, suggesting that despite potential differences between the native Env and the engineered SOSIPs, an allosteric network that was common to both formats was disabled by the mutations.^45^ Indeed, the structure of a PGT151-bound full length Env trimer that was purified by detergent solubilization of cell surface expressed gp160 demonstrated contacts between the topological layers of gp120 and the gp41 three-helix bundle consistent with the conclusion that the mutations were targeting an allosteric network common to both the native Env and engineered SOSIPs.^46^ Recent structures of the full length, detergent-solubilized gp160 SOSIP trimers in complex with PGT145 or PG16 showed a similar overall structure of the SOSIP Env, and also showed similar gp120 topological layer contacts with the gp41 3-helix bundle^47,48^, lending further support for the generality of these contacts and the proposed mechanism of Env allostery. Based on these results, we propose a sequential mechanism by which the HIV-1 Env transitions from the prefusion closed state to a fully open, fusion-competent state through a series of sequential steps (figure 7D). Beginning from a closed state in which the trimer apex and topological layers surround the gp41 three-helix bundle^49^, engagement of a single CD4 induces β20-β21 loop rearrangement and is associated with Env triggering via residue I432 in β20-β21 and residue L193 in V1/V2.^30^ Instability in V1/V2 associated with these changes would allow V3 exposure^34,50^ and initial apex dissociation of a single gp120 protomer from the gp41 three-helix bundle^31^. Internal configurational changes in V1/V2 could then propagate to layers-1 and 2, thereby releasing W571, thus lowering the potential energy barrier to full gp120-gp41 three-helix bundle dissociation resulting in a single gp120 open, asymmetric trimer configuration.^43^ Indeed, viruses with substitution of W571 are replication deficient and non-infectious due to abolished membrane fusion activity despite effective cell surface expression, processing, and retention of CD4 binding, demonstrating the importance of this residue in downstream CD4-induced changes.^51,52^ Based upon our vector analysis (Figure 5D), this transition presumably involves rotation about the W596 anchor toward an transition intermediate. Virologic data is consistent with a role for W596 in mediating transitions, showing mutant dependent differences in syncytia formation and viral entry.^51,53,54^ Absent a stable trimeric apex interface, the remaining two closed state protomers could then decay toward the open state, potentially assisted by CD4 quaternary contacts.^27,49,55^ The observed relationship between the layer-1 and gp41 HR1 C-terminal conformational rearrangement and CD4 triggering suggests W571 displacement with formation of the His-triad may represent an intermediate along the transition path. Previous structural work has implicated the layer-1 His-triad residues in the formation of a second CD4 binding site.^55^ Further, mutation of H66 has been demonstrated in the soluble gp140 SOSIP and membrane anchored gp160 contexts to prevent CD4 induced rearrangements.^10,55,56^ Together, with the mechanistic insights provided by the observed structural rearrangements in the SOSIP Env, the results from this study indicate direct manipulation of the Env allosteric network allows for fine-tuned conformational control of Env structure that can assist in the development and refinement of the next generation of Env vaccine immunogens.

## Methods

### Recombinant HIV-1 envelope SOSIP gp140 production

Antibodies and antibody Fabs were produced as described previously.^57^ BG505 N332 SOSIP gp140 envelopes were expressed as previously described with minor modifications.^58^ Envelope production was performed with Freestyle293 cells (Invitrogen). On the day of transfection, Freestyle293 were diluted to 1.25×10^6^ cells/mL with fresh Freestyle293 media up to 1L total volume. The cells were co-transfected with 650 μg of SOSIP expressing plasmid DNA and 150 μg of furin expressing plasmid DNA complexed with 293Fectin (Invitrogen). On day 6 cell culture supernatants were harvested by centrifugation of the cell culture for 30 min at 3500 rpm. The cell-free supernatant was filtered through a 0.8 μm filter and concentrated to less than 100 mL with a single-use tangential flow filtration cassette and 0.8 μm filtered again. Trimeric Env protein was purified with monoclonal antibody PGT145 affinity chromatography. PGT145-coupled resin was packed into Tricorn column (GE Healthcare) and stored in PBS supplemented with 0.05% sodium azide. Cell-free supernatant was applied to the column at 2 mL/min using an AKTA Pure (GE Healthcare), washed, and protein was eluted off of the column with 3M MgCl_2_. The eluate was immediately diluted in 10 mM Tris pH8, 0.2 μm filtered, and concentrated down to 2 mL for size exclusion chromatography. Size exclusion chromatography was performed with a Superose6 16/600 column (GE Healthcare) in 10 mM Tris pH8, 500 mM NaCl. Fractions containing trimeric HIV-1 Env protein were pooled together, sterile-filtered, snap frozen, and stored at −80 °C.

### Biolayer Interferometry

Cell supernatant mAb binding and dose response curves were obtained using biolayer interferometry (BLI; OctetRed96, FortéBio). Antibodies were immobilized on anti-Human IgG Fc capture (AHC; FortéBio) sensor tips via immersion in 20 μg/ml mAb in phosphate buffered saline (PBS) for 300 s followed by washing in PBS for 60 s at 1000 rpm. For the cell supernatant assays, the mAb captured sensor tips were then immersed in 200 μl of the transfection supernatant for 400 s at 1000 rpm for the association phase after which the sensor tips were immersed in PBS for 600 s for the dissociation phase. The sensor tips were regenerated using glycine pH 2.0 for an immersion time of 20 s between measurements. For the dose response experiments, the sensor tips were immersed in the SOSIP-containing wells for 180 s for the association and 60 s for dissociation phase using a shake speed of 1000 rpm beginning from the lowest SOSIP concentration. Regeneration was not performed between measurements of increase SOSIP concentration. A longer, final dissociation phase of 600 s was measured for the highest concentration SOSIP containing well. Non-specific binding was accounted for via subtraction of sensorgrams obtained using the anti-Flu Hemagglutinin Ab82 control mAb. Reported binding corresponds to values and the end of the association phase. Data was evaluated using the Octet Data Analysis 10.0 software (FortéBio) with dose response curves fitted in GraphPad Prism using the one site specific binding model.

### Surface Plasmon Resonance

Triggering of SOSIP gp140 Envs by soluble CD4 (sCD4) was monitored *via* surface Plasmon resonance and was performed on a BIAcore 3000 instrument (GE Healthcare). Antibodies were immobilized using direct immobilization using amine coupling on CM3 sensor chips (GE Healthcare) at ∼5000 RU. The SOSIP concentration in the presence and absence of a 1:3 mixture of sCD4 was 200 nM. Samples were injected over at a rate of 30 μl/min for a total of 90 μl using the high performance kinject injection mode with a dissociation phase of 600 s over four flow cells containing 17B, 19B, VRC01, or the Ab82 control. The chip surface was regenerated between measurements using two injections of 20 μl of glycine pH 2.0 at a flow rate of 50 μl/min. The resulting response curves were processed using the BIAevaluation 4.1 (GE Healthcare) software using a double reference subtraction. Reported response values were determined taking the average response from 170-175 seconds after the start of the injection.

### Thermal Denaturation

Thermal denaturation experiments were performed using the NanoDSC platform (TA Instruments). Samples were dialyzed into HEPES buffered saline (HBS; 10 mM HEPES, 150 mM NaCl, pH 7.4), diluted in dialysate to 0.2 to 0.4 mg/ml, and degassed for 15 minutes. Following condition of the DSC cells in dialysate, samples were loaded and heated from 20 °C to 100 °C at 3 atm of pressure at a rate of 1 °C/min using the dialysate as a reference. The obtained denaturation profiles were buffer subtracted and base line corrected using a 6^th^-order polynomial using the NanoAnalyze software (TA Instruments). The reported T_max_ corresponds to the average maximum observed heat capacity from three independent measures.

### Cryo-EM Sample Preparation

The BG505 F14 and F14/Vt8 SOSIP trimers complexes were prepared using a stock solution of 2 mg/ml trimer incubated with a six-fold molar excess of VRC01 or VRC03 and 10-1074, respectively. To prevent interaction of the trimer complexes with the air-water interface during vitrification, the samples were incubated in 0.085 mM *n*-dodecyl β-D-maltoside (DDM). Samples were applied to plasma-cleaned QUANTIFOIL holey carbon grids (EMS, R1.2/1.3 Cu 300 mesh) followed by a 30 second adsorption period and blotting with filter paper. The grid was then plunge frozen in liquid ethane using an EM GP2 plunge freezer (Leica, 90-95% relative humidity).

### Cryo-EM Data Collection

Cryo-EM imaging was performed on a FEI Titan Krios microscope (Thermo Fisher Scientific) operated at 300 kV. Data collection images were acquired with a Falcon 3EC Direct Electron Detector operated in counting mode with a calibrated physical pixel size of 1.08 Å with a defocus range between −1.0 and −3.5 µm using the EPU software (Thermo Fisher Scientific). No energy filter or C_s_ corrector was installed on the microscope. The dose rate used was ∼0.8 e^-^/Å^2^·s to ensure operation in the linear range of the detector. The total exposure time was 60 s, and intermediate frames were recorded every 2 s giving an accumulated dose of ∼42 e^-^/Å^2^ and a total of 30 frames per image. A total of 2,350 images for BG505-F14-SOSIP and 2,060 images for BG505-F14/Vt8-SOSIP were collected over two days, respectively.

### Data Processing

Cryo-EM image quality was monitored on-the-fly during data collection using automated processing routines. Initial data processing was performed within cryoSPARC^35^ including particle picking, multiple rounds of 2D classification, *ab initio* reconstruction, homogeneous map refinement and non-uniform map refinement, yielding 3.5 Å and 3.7 Å maps for the BG505-F14-SOSIP and the BG505-F14/Vt8-SOSIP complexes, respectively. Further processing was done outside of cryoSPARC as described next. Movie frame alignment was carried out using UNBLUR^59^, and CTF determination using CTFFIND4^60^. Particles were picked automatically using a Gaussian disk of 90 Å in radius as the search template. For the BG505-F14-SOSIP dataset, 1,518,046 particles were picked from 2,350 micrographs, extracted using a binning factor of 2 and subjected to 8 rounds of refinement in cisTEM^61^, using an *ab-initio* model generated with cryoSPARC^35^. The distribution of scores assigned to each particle by cisTEM showed a clear bi-modal distribution and only particles in the group containing the higher scores were selected for further processing (Supplementary Figure 4). This subset of 77,632 particles was re-extracted without binning and subjected to 10 rounds of local refinement followed by 5 additional rounds using a shape mask generated with EMAN2^62^. Per-particle CTF refinement was then conducted until no further improvement in the FSC curve was observed. At this point, particle frames were re-extracted from the raw data and subjected to per-particle motion correction using all movie frames and applying a data-driven dose weighting scheme as described previously.^63^ The per-particle refinement procedure was iterated two additional times using the newly generated map as a reference and at that point no further improvement was observed. The resolution of the final map was 3.0 Å measured according to the 0.143-cutoff FSC criteria. A b-factor of −120 Å^2^ was applied to the reconstruction for purposes of visualization. For the BG505-F14/Vt8-SOSIP dataset, 869,323 particles were picked from 2,060 micrographs, and a subset of 84,378 particles was used for further local refinement using a similar strategy as that used to process the BG505-F14-SOSIP dataset. The estimated resolution for the final map in this case was 2.9 Å according to the 0.143-cutoff FSC criteria.

### Cryo-EM Structure Fitting

Structure fitting of the cryo-EM maps was performed in Chimera^64^ using the gp120 and gp41 segments from PDB ID 5CEZ (chains G and B, respectively) with F14 and Vt8 mutations added using PyMol. Coordinates for VRC01, VRC03, and 10-1074 were obtained from PDB ID 3NGB (chains H and L), PDB ID 3SE8 (chains H and L), and PDB ID 5T3Z (chains H and L), respectively. Initial coordinate refinement was performed using Rosetta.^65^ The best fit from 110 models was then iteratively with manual refinement in Coot^66^ followed by real-space refinement in Phenix.^67^ Structure fit and map quality statistics were determined using MolProbity^68^ and EMRinger^69^, respectively. Structure and map analyses were performed using a combination of PyMol^70^ and Chimera.

### Vector Based Structure Analysis

Centroids for the vectors in the analysis included a K46-K490 Cα centroid, W571 and W596 c-αs, c-αs of gp120 excluding variable loops the V1/V1 region residues, and the N- and C-termni, and a V1/V2+V3 c-α centroid Vectors between these reference positions were generated and included a projection of the W596 to K46-K490 centroid vector on to the W596 to W571 vector. Angles, distances, and dihedrals between these vectors were then compiled for a set of available crystal and cryo-EM structures with PDB IDs 4TVP^71^, 4ZMJ^25^, 5ACO^72^, 5CEZ^37^, 5CJX^73^, 5D9Q^74^, 5FYJ^75^, 5FYK^75^, 5FYL^75^, 5T3X^76^, 5T3Z^76^, 5U7M^33^, 5U7O^33^, 5UTF^13^, 5V7J^77^, 5V8L^78^, 5V8M^78^, 6CDE^38^, 6CDI^38^, 6CH7^79^, 6CH8^79^, 6CUE^80^, 6CUF^81^, and 6DE7^12^, 5THR^26^, 5VN3^27^, 5VN8^73^, 6CM3^31^, 6EDU^31^, 5FUU^82^, 6DCQ^46^, 6MAR^83^, 5U1F^55^, 5C7K^84^, 5I8H^85^, 5UM8^86^, 5UTY^13^, 5VIY^87^, 5VJ6^87^, 5WDU^14^, 6B0N^88^, 6CCB^89^, 6CE0^11^, 6CH9^79^, 6CHB^79^, 6CK9^90^, 6DCQ^46^, 6DID^91^, 6E5P^92^, 6IEQ^93^, 6MCO^94^, 6MDT^94^, 6MN7^95^, 6MPG^96^, 6MPH^96^, 6MTJ^97^, 6MTN^97^, 6MU6^97^, 6MU7^97^, 6MU8^97^, 6MUF^97^, 6MUG^97^, 6N1V^96^, 6N1W^96^, 6NC2^98^, 6NC3^98^, 6NF2^96^, 6NIJ^47^, 6NM6^99^, 6NNF^99^, 6NNJ^99^, 6OHY^100^, 6OKP^101^, 6OLP^47^, 6ORN^39^, 6ORO^39^, 6ORP^39^, 6ORQ^39^, 6OSY^96^, 6OT1^96^, and 6UDA^40^ as well as the BG505 F14 and F14/Vt8 SOSIP structures determined in this study. Vector based structural analysis was performed using the VMD^102^ Tcl interface. Principal component analysis of the resulting vectors, angles, and torsions was performed using R with the data centered and scaled^103^.

### Cell-Surface Expressed Env gp160 Antigenicity

FreeStyle 293F cells (ThermoFisher, catalog #R79007) (1 x 10^6^ cells/ml, 0.5ml per well of a 12-well plate) were transfected using a mixture of 1μg HIV Env gp160s DNA in 75 μl jetPRIME buffer (Polyplus-transfection) with 2 μl jetPRIME transfection reagent (Polyplus-transfection) following the manufacturer’s protocol. Transfected cells were cultured in Freestyle 293 Expression Medium (Invitrogen Inc.) at 120rpm for 48 hours before flow cytometry staining. 293F cells were counted using trypan blue and then were rinsed with PBS containing 1% BSA, pelleted at 250 × g for 3 min. Cells were incubated with wither sCD4 or eCD4-Ig^41^ at a final concentration of 10 μg/mL for 20 min at 4 °C. Human anti-HIV Env Abs at a final concentration of 10 μg/mL were used to stain 0.4 × 10^6^ cells per well in 40 μL PBS containing 1% BSA of in V bottom 96-well plates, 30 min at RT in the dark. Cells were then washed once with 200 μL PBS containing 1% BSA and incubated with PE conjugated Goat F(ab’)2 Anti-Human IgG-(Fab’)2 secondary antibody (abcam, Cambridge, MA) at a final concentration of 2.5 μg/ml in 100 μL PBS containing 1% BSA per well. After 30 min incubation at 4 °C in the dark, cells were washed once, and stained with 200 μL aqua viability dye (1:1000 in PBS) for 20 min at RT in the dark, then washed twice with PBS containing 1% BSA. Flow cytometric data were acquired on a LSRII using FACSDIVA software (BD Biosciences) and were analyzed with FlowJo software (FlowJo). FACSDIVA software (BD Biosciences) and were analyzed with FlowJo software.

### Preparation and smFRET analysis of dye-labeled HIV-1_BG505_ virus Env

Dye-labeled wild-type and mutant F14Vt8 HIV-1_BG505_ virus Env were prepared and imaged as previously described.^45^ Briefly, peptides-tagged Envs in the context of full-length virus were constructed by introducing two labeling tag peptides (GQQQLG; GDSLDMLEWSLM) into variable loops V1 and V4 of gp120 subunit on both wild-type and F14Vt8 BG505 Env in the context of a replication competent clade A virus carrying the Q23 as the backbone. Wild-type and mutant BG505 virus Env for smFRET imaging was prepared by co-transfecting HEK293 cells with mixed plasmids consisting of a 40:1 ratio of replication-incompetent (RT deleted) wild-type vs. peptides-tagged HIV-1_BG505_ virus Env. The viruses were harvested 40-hour post-transfection, filtered, concentrated, and labeled with donor dye Cy3B(3S)-cadaverine (0.5 μM) and acceptor dye LD650-CoA (0.5 μM, Lumidyne Technologies) in the presence of transglutaminase (0.65 μM, Sigma Aldrich) and AcpS (5 μM) at room temperature overnight. PEG2000-biotin (0.02 mg/ml, Avanti Polar Lipids) was then added to above labeling solutions the next day and incubated for 30 minutes prior to purification. Labeled viruses were further purified by ultracentrifugation at 40,000 rpm over a 6%-18% Optiprep (Sigma Aldrich) gradient for 1 hour.

Dye-labeled virus Envs were immobilized on a streptavidin-coated quartz microscopy slide for smFRET imaging. smFRET movies were acquired on a prism-based total internal reflection fluorescence microscope, and FRET traces were extracted and analyzed by using a customized software package based on LabView and Mathworks.^104^ The evanescent field generated by a 532 nm CW laser (Opus, Laser Quantum) was used to excite donor dye. Fluorescence from both donor and acceptor dyes that are labeled on HIV-1_BG505_ virus Env was firstly collected, then spectrally split, and later simultaneously detected by two synchronized sCMOS cameras (ORCA-Flash4.0v2, Hamamatsu). The dynamics of fluorescently labeled Env on virus was traced by monitoring fluorescence signals with 40 milliseconds time-resolution for 80 seconds in imaging buffer containing 50 mM Tris pH 7.4, 50 mM NaCl, a cocktail of triplet-state quenchers, and oxygen scavenger consisted of 2 mM protocatechuic acid with 8 nM protocatechuic-3,4-dioxygenase. In the case of ligands binding experiments, fluorescently labeled wild-type or F14Vt8 mutant virus Env was pre-incubated with 10 μg/ml sCD4D1D2– Igαtp (12xCD4) for 30 min at room temperature before imaging.

FRET (presented as efficiency/value) traces in real time were derived from fluorescence signals traces based on the equation FRET= I_A_/(γI_D_+I_A_), where I_D_ and I_A_ are the fluorescence intensities of donor (I_D_) and acceptor (I_A_), and γ is the correlation ecoefficiency. FRET traces that pass through the stringent filters of signal-to-noise (S/N) ratio as well as anti-correlated features between I_D_ and I_A_ were used to build FRET histograms. On the basis of the observations from original FRET traces and the idealization by hidden Markov modeling, FRET histograms were further fitted into the sum of three Gaussian distributions. Relative state occupancy of each FRET state was estimated as the area under each Gaussian curve, where standard errors was calculated from thousands of individual data points that corresponds to the indicated number (N) of molecules in FRET histogram. The smFRET approach has been extensively validated as described in Lu et al. Specifically, the fluorophores applied are not observed to exhibit abnormal dye behavior or photo-physical effects. smFRET data for labeled HIV-1 Env with positions of donor and acceptor fluorophores reversed, yields identical results. Similarly, identical results are observed when fluorophores are attached to Env using amino acids rather than peptide epitope tags. Last but not least, monitoring conformational dynamics of Env from a different perspective identifies the same three conformational states.^45^

## Acknowledgements

This work was supported by NIAID Division of AIDS Center for HIV/AIDS Vaccine Immunology and Immunogen Discovery (CHAVI-ID, grant number UM1-AI100645, B.F.H.), NIAID extramural project grant 1R01AI145687-01 (P.A.), NIGMS extramural project grant R01-AI150560 (W.M., S.C.B.), NIAID grant R01-GM098859 (S.C.B), start-up funding from the Translating Duke Health Initiative and the Duke University Center for AIDS Research (CFAR), an NIH funded program (5P30 AI064518) (P.A.), the US National Institutes of Health Intramural Research Program, US National Institute of Environmental Health Sciences (ZIC ES103326 to M.J.B), and the NIH Ruth L. Kirschstein National Research Service Award (NRSA) 5T32AI007392 (R.H). Cryo-EM data were collected at the Shared Materials Instrumentation Facility at Duke University. This work utilized the Duke Compute Cluster and benefited from the Duke Data Commons storage supported by the NIH (1S10OD018164-01). The authors thank John Mascola and James Arthos for providing the eCD4-Ig and 12xCD4 reagents, respectively.

## Author Contributions

R.H. conceived and designed the mutations, performed experiments, wrote and edited the manuscript. RH, BFH and SMA conceived the idea and implementation of the research studies. K.S., K.W., B.F.H., P.A. and S.M.A. edited the manuscript. R.H., and S.M.A. designed the research studies involving mutant selection, biophysical and antigenicity characterization. R.H. and P.A. designed the cryo-EM studies. R.H. and K.W. designed mutant Env panel. K.S. produced and purified antibodies and produced Env proteins. R.H and K.S. purified Env proteins. R.H. performed and analyzed transfection supernatant BLI. E.C. performed SPR and dose response BLI and analyzed data. R.J.E. performed negative stain-EM and analyzed data. R.H. performed DSC and analyzed data. R.H., Y.Z., A.B., and P.A. determined cryo-EM structures. R.H. and P.A. performed structural analysis. A.H. and M.J.B collected cryo-EM data. R.P. and Q.H. performed cell surface binding experiments and analyzed data. M.L., W.M., and S.B. performed smFRET and analyzed data. All authors read and approved manuscript.

## Data and Code Availability Statement

The datasets and code generated and/or analyzed during this study are available from the corresponding author on reasonable request. The cryo-EM structures are being deposited to the EMDB.

