## Supplemental Data for "Disruption of the HIV-1 Envelope allosteric network blocks CD4-induced rearrangements"

### Supplementary Data and Figures

Rory Henderson[1,4,*,#], Maolin Lu[6], Ye Zhou[8], Zekun Mu[11], Robert Parks[4], Qifeng Han[1,4], Allen L. Hsu[5], Elizabeth Carter[1,4], Scott C. Blanchard[7&], RJ Edwards[1,4], Kevin Wiehe[1,4], Kevin O. Saunders[2,4], Mario J. Borgnia[5], Alberto Bartesaghi[8,9,10], Walther Mothes[6], Barton F. Haynes[1,4], Priyamvada Acharya[2,4,*], S. Munir Alam[1,3,4,*]

Supplementary Table 1: BG505 SOSIP Env Mutations

| Mutation | Mutations |
|---|---|
| *F1* | V68I, S115V, A204L, V208L, V255W, N377L, M426W, M434W, H66S |
| *F2* | V68I, S115V, A204L, V208L, V255W, H66S |
| *F3* | V68I, A204V, V208L, V255L, H66S |
| *F4* | V68I, S115V, V208L, V255L, H66S |
| *F5* | V68I, S115V, A204L, V208L, V255W, N377L, H66S |
| *F6* | V68I, S115V, A204L, V208L, V255L, W69L |
| *F7* | V68I, S115V, A204L, V208L, V255L, W69V |
| *F8* | V68I, S115V, A204L, V255L, V208L, W69A |
| *F9* | V68I, S115V, A204L, V208L, V255W, N377L, M426W, H66S |
| *F10* | V68I, S115V, A204L, V208L, V255W, N377L, M434W, H66S |
| *F11* | V68I, S115V, A204V, V208L, V255L, H72P, H66S |
| *F12* | V68I, S115V, V208L, V255L, H66K |
| *F13* | V68I, S115V, A204L, V208L, V255W, N377L, M426W, M434W |
| *F14* | V68I, A204V, V208L, V255L |
| *F15* | V68I, A204L, V208L, V255W, N377L |
| *Vt1* | Y177F, T320L, D180A, Q422L, Y435F, Q203M, E381L, R298M, N302L, N300L |
| *Vt2* | Y177F, T320L, D180A, Q422L, Y435F, Q203M, N302L, N300L |
| *Vt3* | Y177F, T320L, D180A, Q422L, Y435F, Q203M, R298M, N302L, N300L |
| *Vt4* | Y177F, T320L, D180A, Q422L, Y435F, Q203M, E381L, N302L, N300L |
| *Vt5* | T320L, D180A, Q422L, Q203M, E381L, R298M, N302L, N300L |
| *Vt6* | T320L, D180A, Q422L, Q203M, N302L, N300L |
| *Vt7** | I201C, A433C, L154M, N300M, N302M, T320L |
| *Vt8* | T320M, Q422M, Q203M, N302L, N300L |
| *Vt9* | T320M, Q422M, Q203M, N302L, N300L, S174V |

* Design DS-SOSIP.4mut from [13]

#### Supplementary Table 2: CD4 Binding Kinetics and Affinities

| CD4 Binding* | $k_a$ ($M^{-1}s^{-1}$) | $k_d$ ($s^{-1}$) | $K_d$ (nM) |
|---|---|---|---|
| **BG505 WT** | 1.12E+04 ± 5.77E+03 | 6.10E-04 ± 9.50E-05 | 67.9 ± 26.4 |
| **BG505 F14** | 1.72E+04 ± 4.75E+03 | 1.13E-03 ± 1.00E-04 | 73.0 ± 26.2 |
| **BG505 Vt8** | 9.44E+03 ± 6.65E+02 | 7.81E-04 ± 6.40E-05 | 83.9 ± 12.6 |
| **BG505 F14/Vt8** | 2.92E+04 ± 5.35E+03 | 4.52E-04 ± 7.40E-05 | 15.6 ± 0.4 |

* Values reported as the average of two replicate measures ± one standard deviation

#### Supplementary Table 3: bnAb Binding Affinities ($K_d$ [nM])

| | BG505 WT | BG505 F14 | BG505 Vt8 | BG505 F14/Vt8 |
|---|---|---|---|---|
| **PGT145** | 64.2 | 76.52 | 68.04 | 80.2 |
| **PGT151** | 26.9 | 23.34 | 24.04 | 24.04 |
| **VRC01** | 611.7 | 565.7 | 268.9 | 313.7 |
| **PG9** | 444 | 45.9 | 335 | 336.8 |
| **VRC26** | 67.6 | 50.23 | 60.4 | 50.6 |
| **PGT121** | 222.3 | 218.6 | 260.3 | 262 |

* Values reported from fit to average the average value of two experiments

Supplementary Table S4: Cryo-EM Data Collection and Refinement Statistics

| | BG505 F14 SOSIP | BG505 F14/Vt8 SOSIP |
|---|---|---|
| *Data Collection* | | |
| Microscope | FEI Titan Krios | FEI Titan Krios |
| Voltage (kV) | 300 | 300 |
| Electron dose ($e^-/Å^2$) | 42 | 42 |
| Detector | Falcon 3 | Falcon 3 |
| Pixel Size (Å) | 1.08 | 1.08 |
| Defocus Range (µm) | ~1.5-3 | ~1.5-3 |
| Magnification | 75000 | 75000 |
| *Reconstruction* | | |
| Software | cisTEM | cisTEM |
| Particles | 77632 | 84378 |
| Symmetry | C3 | C3 |
| Box size (pix) | 320 | 320 |
| Resolution (Å) (FSC0.143)* | 3.0 | 2.9 |
| *Refinement (Phenix)* | | |
| Protein residues | 999 | 1472 |
| Chimera CC | 0.77 | 0.63 |
| R.m.s. deviations | | |
| Bond lengths (Å) | 0.01 | 0.01 |
| Bond angles (°) | 1.06 | 1.24 |
| *Validation* | | |
| Molprobity score | 1.71 | 1.87 |
| Clash score | 4.4 | 6.0 |
| Favored rotamers (%) | 99.4 | 97.2 |
| Ramachandran | | |
| Favored regions (%) | 91.6 | 90.3 |
| Disallowed regions (%) | 0.41 | 0.6 |

* Resolutions are reported according to the FSC 0.143 gold-standard criterion

Supplementary Table S5: PCA Cluster PDBs

| **Red** | **Pink** | **Green** | **Blue** | **Cyan** | **Black** |
|---|---|---|---|---|---|
| 6b0n | 5u7o | 6okp | 6osy | *F14* | *F14Vt8* |
| 5fyj | 4zmj | 5um8 | 6n1w | 6n1v | 6uda |
| 6nnf | 6mco | 5cjx | 6nc3 | 6orn | 6orq |
| 5v8m | 6muf | | 6nf2 | | 6orp |
| 6ck9 | 5t3z | | 6cdi | | 6oro |
| 5fyk | 6mug | | 6mph | | |
| 6nm6 | 5d9q | | 6mpg | | |
| 6mtn | 6ch8 | | 6ot1 | | |
| 5cez | 4tvp | | 6cde | | |
| 6nnj | 5i8h | | 6cuf | | |
| 6de7 | 5utf | | 6cue | | |
| 5aco | 5t3x | | | | |
| 6ohy | 6ch7 | | | | |
| 6ieq | | | | | |
| 6mtj | | | | | |
| 6mu7 | | | | | |
| 6mu8 | | | | | |
| 6mu6 | | | | | |
| 5v7j | | | | | |
| 5v8l | | | | | |
| 6mdt | | | | | |
| 5fyl | | | | | |
| 5uty | | | | | |
| 5u7m | | | | | |

Supplementary Table 6: Cell Surface Expressed Trimer - Percentage of Cells Binding

| | Control | BG505 | BG505 DS | BG505 F14/Vt8 | BG505 F14 | BG505 Vt8 |
|---|---|---|---|---|---|---|
| **N6** | 0.3 ± 0.2 | 45.3 ± 8.1 | 41.0 ± 5.8 | 40.3 ± 3.7 | 53.9 ± 12.0 | 44.4 ± 5.0 |
| **CH01** | 0.2 ± 0.1 | 32.8 ± 2.6 | 20.8 ± 9.7 | 22.1 ± 11.9 | 37.7 ± 12.4 | 30.2 ± 3.0 |
| **PGT125** | 0.2 ± 0.1 | 47.1 ± 6.6 | 38.2 ± 9.1 | 37.9 ± 5.2 | 54.7 ± 10.1 | 43.2 ± 10.5 |
| **PGT145** | 0.3 ± 0.2 | 46.0 ± 4.6 | 35.4 ± 3.0 | 34.8 ± 2.6 | 48.8 ± 11.9 | 38.2 ± 4.5 |
| **17B** | 0.3 ± 0.1 | 9.0 ± 3.6 | 0.6 ± 0.2 | 1.2 ± 0.9 | 2.7 ± 1.0 | 0.8 ± 0.5 |
| **19B** | 0.2 ± 0.2 | 17.1 ± 8.2 | 16.8 ± 1.7 | 8.0 ± 4.8 | 23.5 ± 8.4 | 8.0 ± 5.4 |
| **17B + sCD4** | 0.2 ± 0.1 | 25.9 ± 7.1 | 1.3 ± 0.2 | 0.7 ± 0.3 | 3.7 ± 1.2 | 0.9 ± 0.5 |
| **19B + sCD4** | 0.2 ± 0.1 | 25.3 ± 8.9 | 16.9 ± 8.3 | 9.0 ± 5.5 | 23.6 ± 8.0 | 9.0 ± 6.4 |
| **17B + CD4Ig** | 0.2 ± 0.1 | 32.9 ± 6.5 | 0.9 ± 0.3 | 0.8 ± 0.6 | 8.2 ± 0.8 | 9.5 ± 1.1 |
| **19B + CD4Ig** | 0.3 ± 0.2 | 38.3 ± 8.2 | 15.0 ± 7.1 | 9.5 ± 5.6 | 27.0 ± 7.1 | 21.8 ± 5.5 |

* Values reported as the average of three experiments with standard deviations

Supplementary Table 7: Cell Surface Expressed Trimer - MFI

| | Control | BG505 | BG505 DS | BG505 F14/Vt8 | BG505 F14 | BG505 Vt8 |
|---|---|---|---|---|---|---|
| **N6** | 39.9 ± 14.1 | 457.3 ± 137.9 | 338.0 ± 72.6 | 341.3 ± 43.2 | 568.0 ± 152.6 | 378.3 ± 73.1 |
| **17B** | 34.1 ± 11.5 | 103.4 ± 38.6 | 49.2 ± 24.2 | 51.6 ± 28.0 | 60.0 ± 29.0 | 51.0 ± 25.0 |
| **19B** | 33.1 ± 11.5 | 211.3 ± 73.2 | 198.3 ± 67.9 | 109.3 ± 39.0 | 265.0 ± 84.2 | 103.8 ± 45.4 |
| **17B + sCD4** | 33.7 ± 11.1 | 244.7 ± 65.0 | 51.6 ± 27.1 | 48.8 ± 24.1 | 67.8 ± 34.0 | 48.4 ± 23.4 |
| **19B + sCD4** | 33.5 ± 11.4 | 306.7 ± 83.0 | 194.0 ± 63.0 | 115.2 ± 37.7 | 255.0 ± 88.4 | 114.8 ± 53.6 |
| **17B + CD4Ig** | 42.5 ± 25.1 | 400.0 ± 115.2 | 50.2 ± 23.9 | 50.3 ± 28.4 | 102.4 ± 28.9 | 90.8 ± 29.1 |
| **19B + CD4Ig** | 35.5 ± 13.1 | 531.3 ± 134.1 | 176.3 ± 62.7 | 119.5 ± 45.9 | 269.7 ± 63.2 | 190.7 ± 43.1 |

* Values reported as the average of three experiments with standard deviations

Supplementary Table 8: smFRET Statistics

| gp160 | BG505 WT | BG505 WT + 12xCD4 | BG505 F14/Vt8 | BG505 F14/Vt8 + 12xCD4 |
|---|---|---|---|---|
| **State 1** | 46 ± 7 | 20 ± 8 | 49 ± 9 | 43 ± 7 |
| **State 2** | 26 ± 8 | 30 ± 11 | 20 ± 9 | 22 ± 11 |
| **State 3** | 28 ± 10 | 50 ± 13 | 31 ± 12 | 35 ± 12 |

**Supplementary Figure 1**

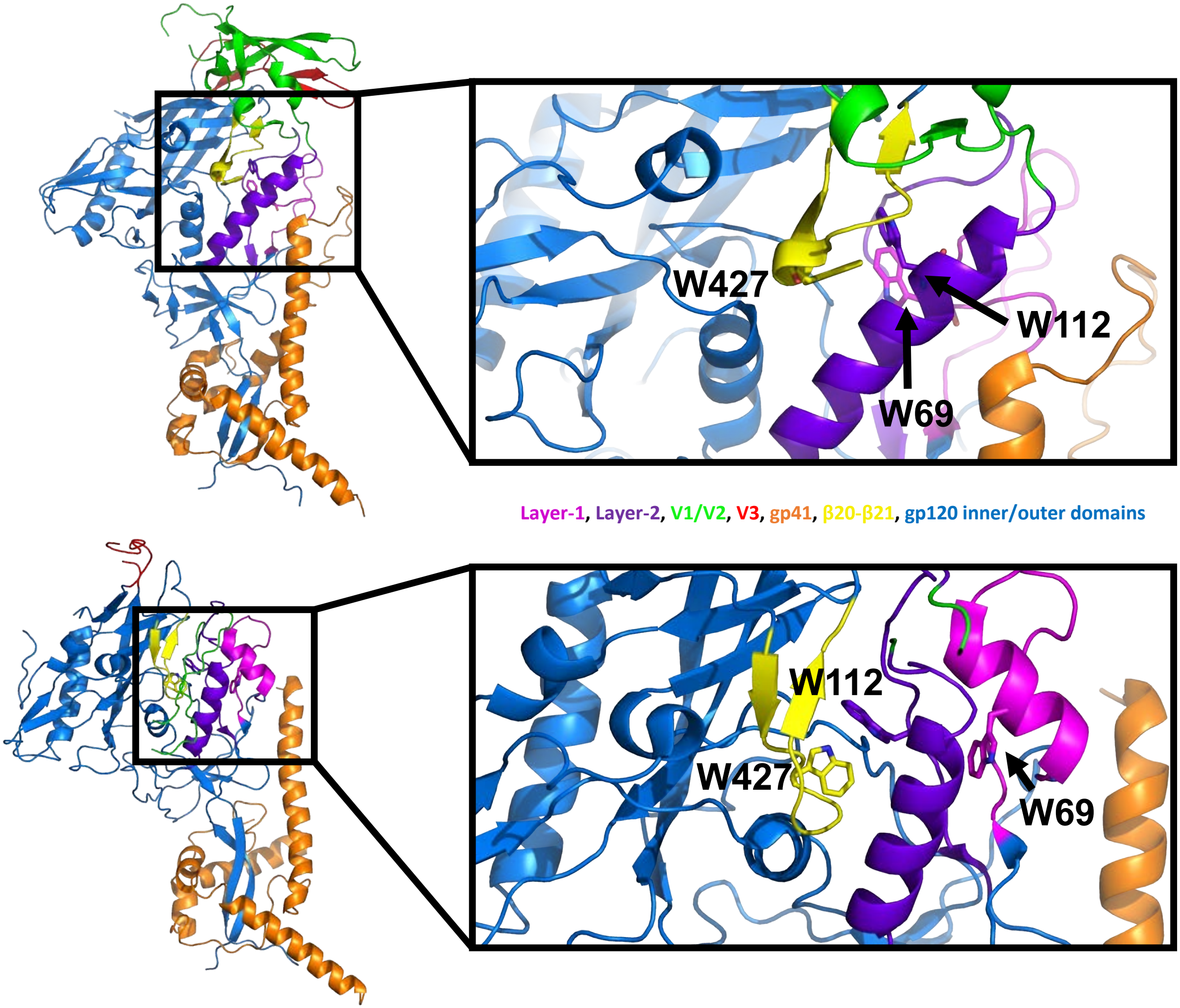


**Supplementary Figure 1.** *Structure of a closed and open state HIV-1 Env gp140 SOSIP protomer.* ***(A) (left)*** A closed state SOSIP gp140 protomer (PDB ID 5CEZ) identifying key structural elements. ***(right)*** Elements of the closed state allosteric network including β20-β21 (yellow), layer-1/2 (lime green/purple), V1/2/3 (green/red) and tryptophans 69, 112 and 427. ***(B) (left)*** An open state SOSIP gp140 protomer (PDB ID 5VN3) identifying key structural elements. ***(right)*** Elements of the open state allosteric network including β20-β21 (yellow), layer-1/2 (lime green/purple), V1/2/3 (green/red) and tryptophans 69, 112 and 427.

### Supplementary Figure 2

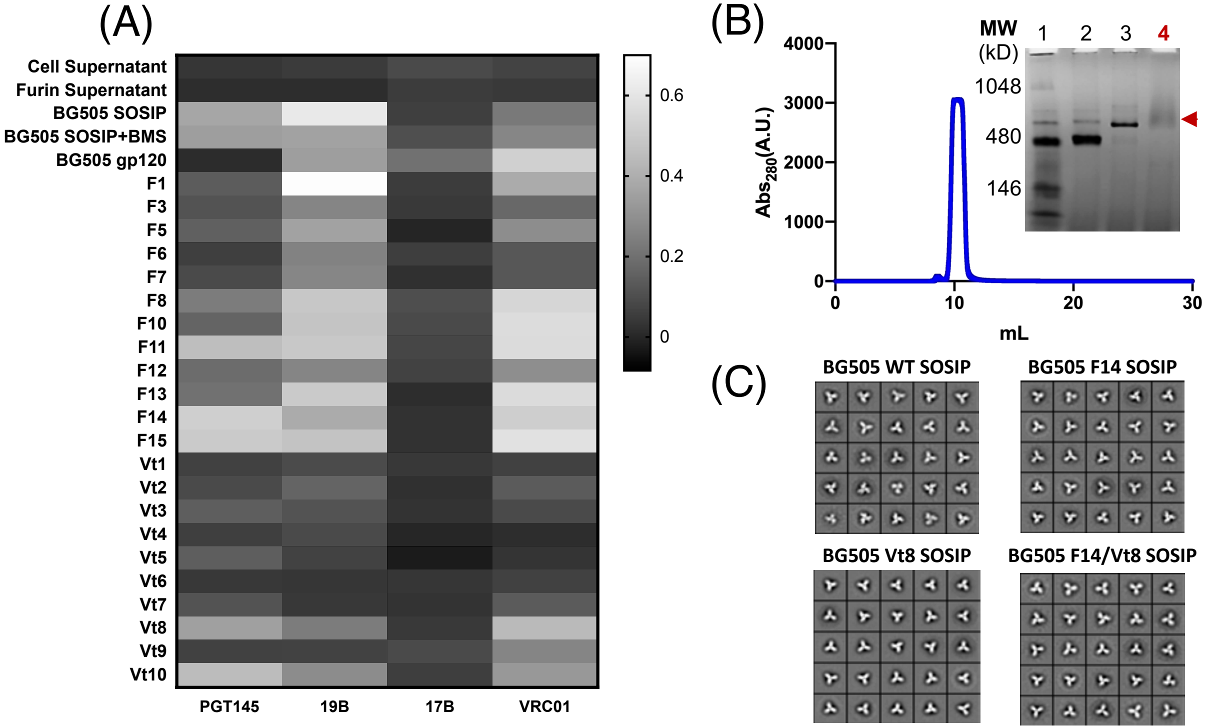


**Supplementary Figure 2.** *BG505 SOSIP Mutant Screening Results* ***(A)*** Transfection cell supernatant BLI heatmap for F-series, Vt-series, WT SOSIP, WT gp120, BMS-626529-bound BG505 SOSIP, and controls binding to PGT145, 19B, 17B, and VRC01. Results plotted as mean response (nm) from triplicate transfection supernatant screening. ***(B)*** *(graph)* Representative size exclusion chromatogram from the BG505 F14 SOSIP purification post PGT145 affinity column purification. *(inset)* Representative non-reducing SDS-PAGE gel for the BG505 SOSIP.664 mutants. Lanes 1-3 include protein marker, Ferritin, and Thyroglobulin, respectively. Lane 4 contains the SOSIP trimer. ***(C)*** Negative-stain EM two-dimensional class averages for purified BG505 WT, F14, Vt8, and F14/Vt8 SOSIPs.

#### Supplementary Figure 3

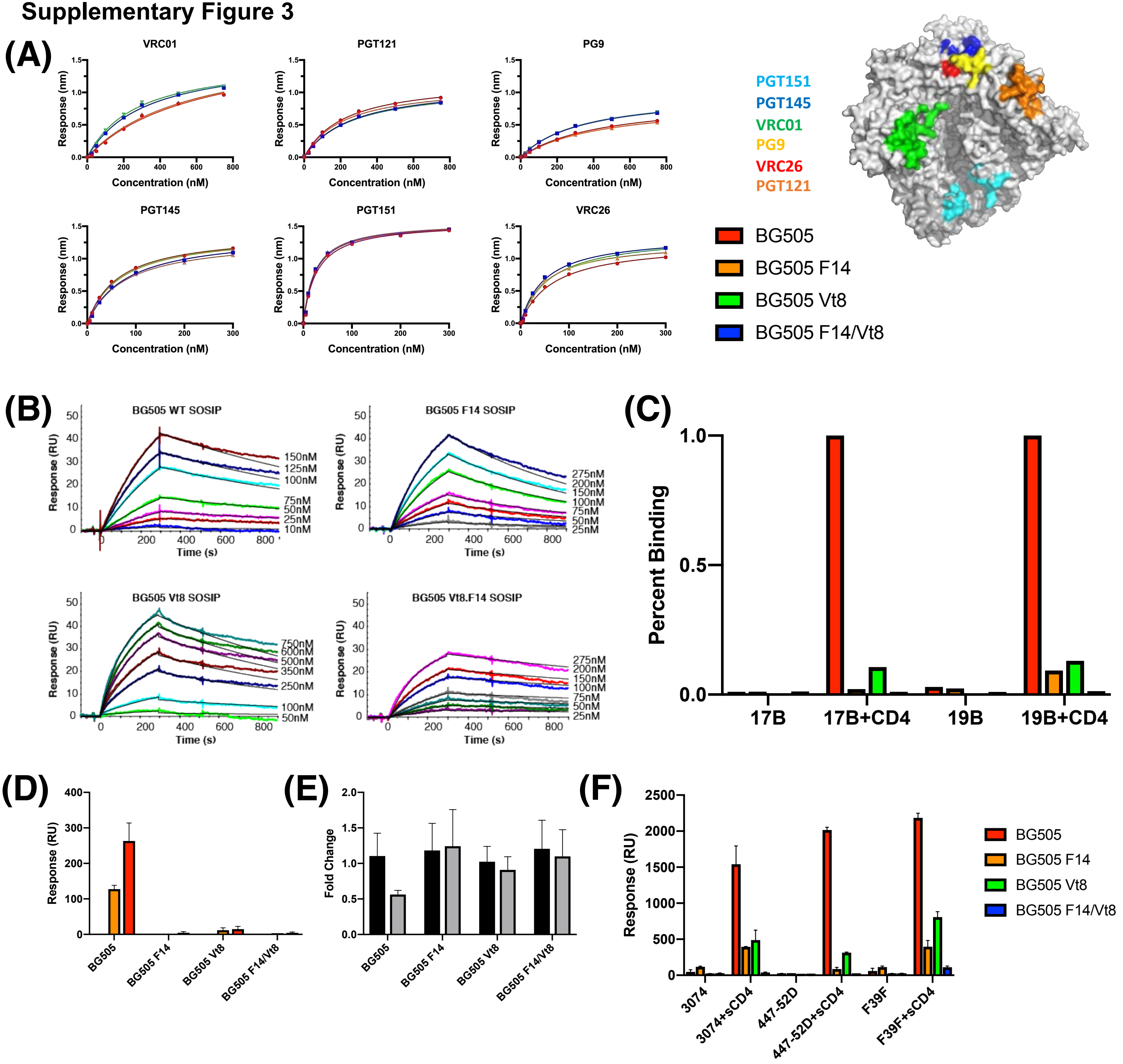

**Supplemental Figure 3.** *BG505 SOSIP Mutant bnAb Binding, CD4 binding, and CD4 Triggering* ***(A)*** *(left)* Dose response curves for binding of VRC01, PGT121, PG9, PGT145, PGT151, and VRC26 to the BG505 WT, F14, Vt8, and F14/Vt8 SOSIPs. Results plotted as mean response (nm) from duplicate experiments (mean standard error is within plotted points). Lines represent one-site specific fitting of BLI binding data in Prism. (*right*) Surface representation of BG505 SOSIP with bnAb epitopes highlighted. ***(B)*** Representative SPR titrations for BG505 WT, F14, Vt8, and F14/Vt8 SOSIP binding to CD4-Ig. ***(C)*** Representative CD4 triggering results for BG505 WT (red), F14 (orange), Vt8 (Vt8), and F14/Vt8 (blue) SOSIP. Responses are normalized to BG505 WT SOSIP. ***(D)*** Response values for 17B interaction with BG505 WT, F14, Vt8, and F14/Vt8 SOSIP alone (black) or incubated with sCD4 for either 30 minutes (orange) or 20 hours (red). Data are for two measurements. ***E)*** Response ratio for PGT145 interaction with BG505 WT, F14, Vt8, and F14/Vt8 SOSIP incubated with sCD4 for either 30 minutes (black) or 20 hours (grey) relative to the relevant unliganded trimer. Data are for two measurements. ***F)*** Response values for V3 targeting 3074, 447-52D, and F39F interaction with BG505 WT, F14, Vt8, and F14/Vt8 SOSIP in the absence and presence of sCD4.

Supplementary Figure 4

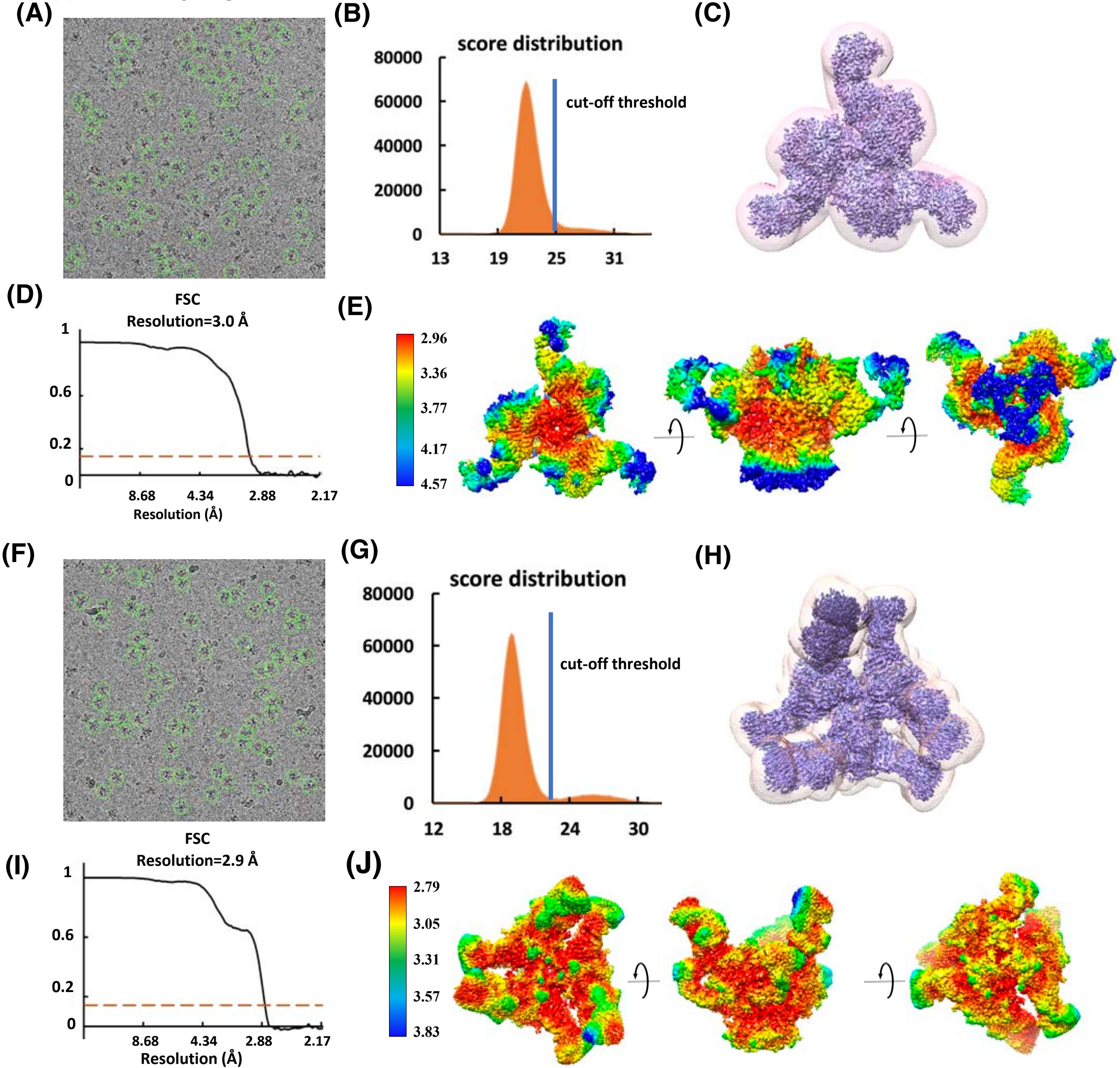

**Supplementary Figure 4**

**Supplementary Figure 4.** *Cryo-EM data analysis for BG505 F14 and F14/Vt8 SOSIP datasets.* **(A,F)** Representative micrographs showing particles selected for refinement from BG505-F14-SOSIP **(A)**, and BG505-F14/Vt8-SOSIP **(F)** datasets. **(B,G)** Distribution of cisTEM score values assigned to particles extracted from datasets BG505-F14-SOSIP **(B)**, and BG505-F14/Vt8-SOSIP **(G)**. **(C,H)** Cryo-EM map (solid) with superimposed shape mask (transparent) used during 3D refinement for BG505-F14-SOSIP **(C)**, and BG505-F14/Vt8-SOSIP **(H)** datasets. **(D,I)** Fourier Shell Correlation curves between half-maps showing estimated resolution according to the 0.143-cutoff criteria (dashed line) for BG505-F14-SOSIP **(D)** and BG505-F14/Vt8-SOSIP **(I)** reconstructions. **(E,J)** Local map resolution determined using RELION-3.0 for reconstructions of BG505-F14-SOSIP **(E)** and BG505-F14/Vt8-SOSIP **(J)**.

#### Supplementary Figure 5

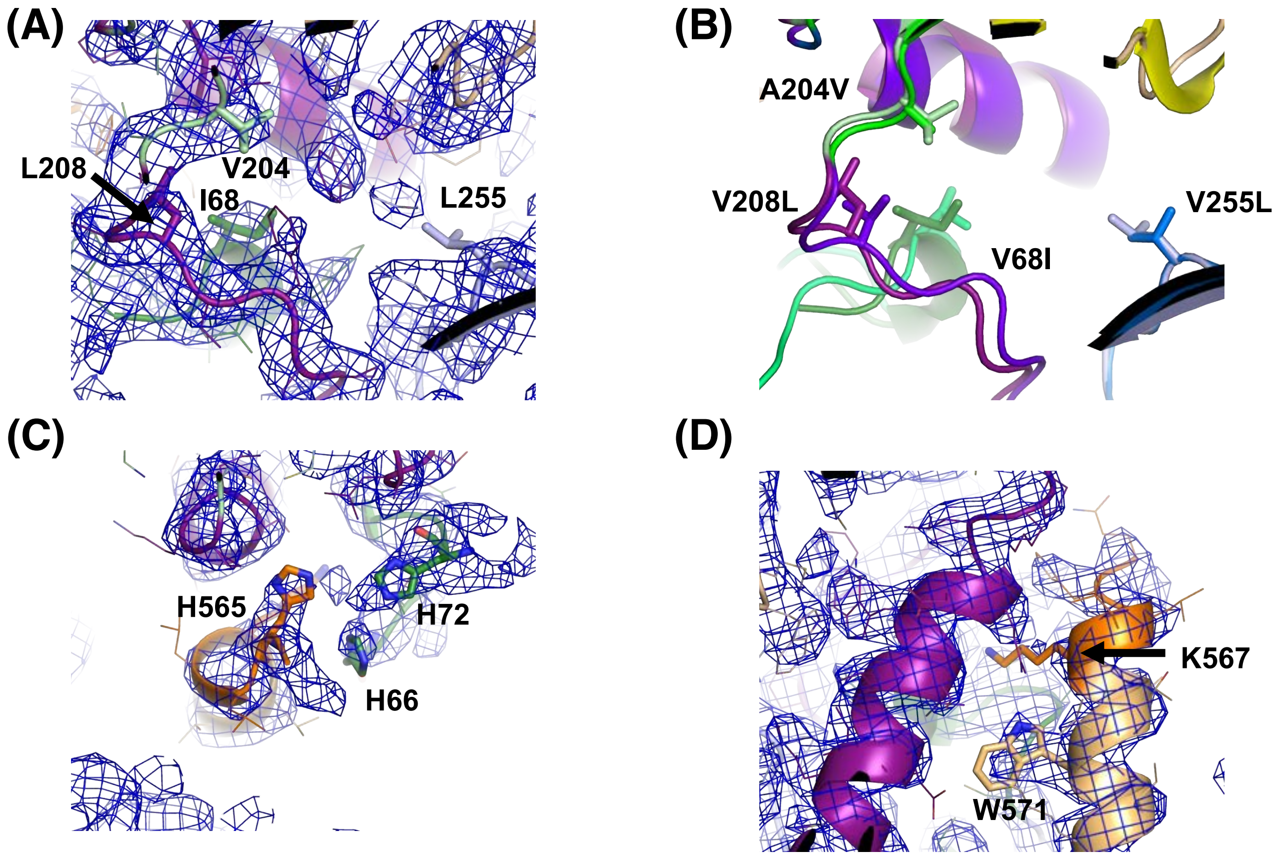


Layer-1, Layer-2, V1/V2, V3, gp41, B20-β21, gp120 inner/outer domains

**Supplementary Figure 5.** *Cryo-EM fit of BG505 F14 mutations.* **(A)** Cryo-EM map of the BG505 F14 construct with fitted coordinates depicting the F14 mutation sites. **(B)** Alignment of the BG505 F14 SOSIP gp120 with the BG505 WT SOSIP (PDB ID 5CEZ) gp120 showing the relative positions of the F14 mutations. ***(C)*** Cryo-EM map (mesh) depicting H66, H72, and H565. ***(D)*** Cryo-EM map (mesh) depicting position of gp41 residues K567 and W571.

#### Supplementary Figure 6

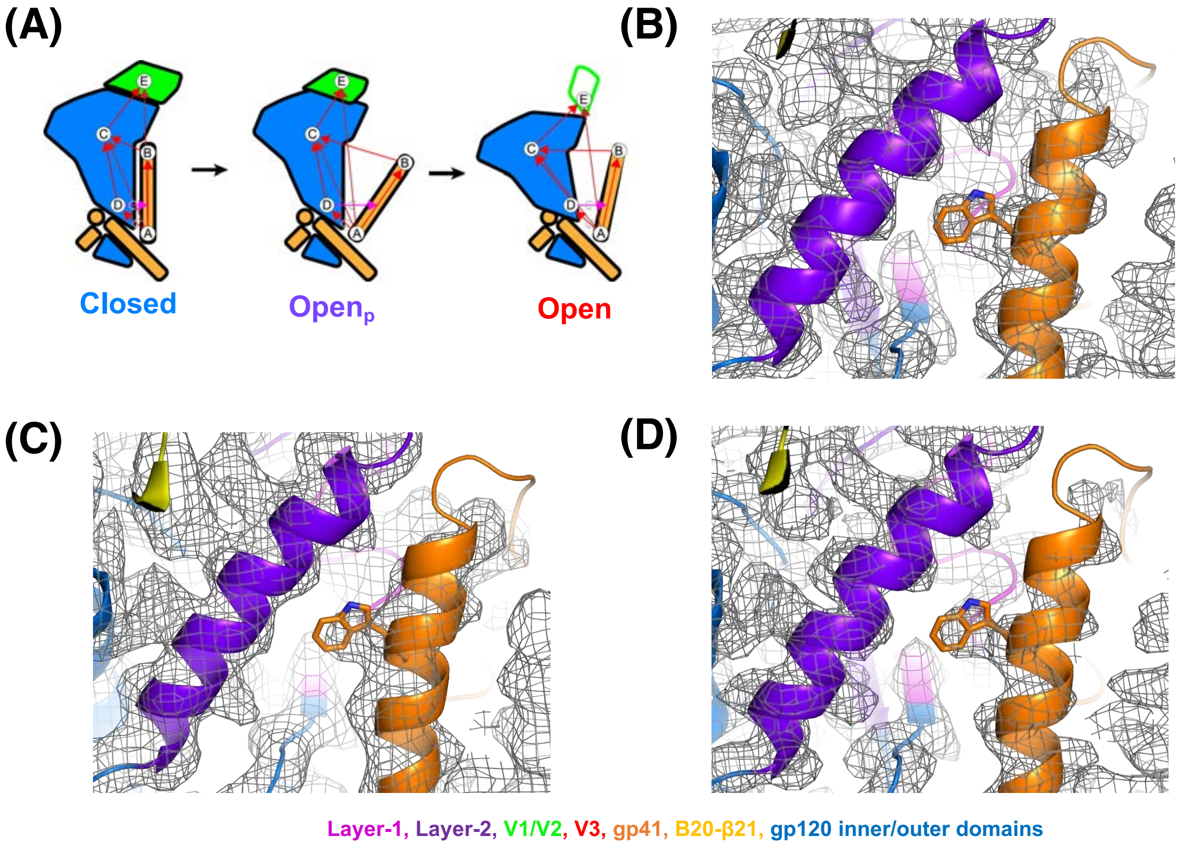


**Supplementary Figure 6.** *Vectors based analysis and cryo-EM fit of BG505 F14/Vt8 SOSIP coordinates to vFP bound BG505 DS SOSIP cryo-EM maps.* ***(A)*** Cartoon representations of gp120 (blue), gp120 V1/V2 region (green), and gp41 (yellow) in the closed, partially open, and open states. Letters indicate location of centroids with arrows depicting vectors between centroids. The reduced size of the V1/V2 region in the partially open state represents rearrangement in this region while the green outline of V1/V2 in the open state indicates complete dissociation and disorder. ***(B)*** Fit of the BG505 F14/Vt8 coordinates into the vFP20.01 bound BG505 DS SOSIP cryo-EM map (EMD-*7459)* ***(C)*** Fit of the BG505 F14/Vt8 coordinates into the vFP16.02 bound BG505 DS SOSIP cryo-EM map (EMD-*7460)* ***(D)*** Fit of the BG505 F14/Vt8 coordinates into the vFP1.01 bound BG505 DS SOSIP cryo-EM map (EMD-*7622)*

**Supplementary Figure 7**

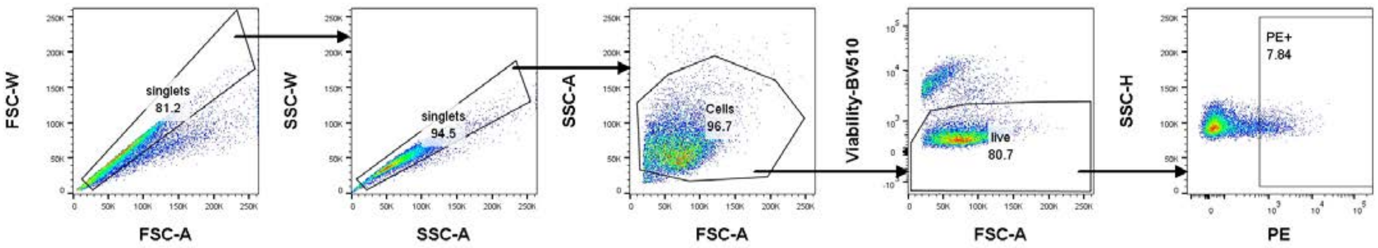


**Supplementary Figure 7.** ***F****low cytometry gating strategy.* Example data depicting the gating strategy used to assess Env cell surface expressed gp160 trimer binding to bnAbs and non-bnAbs.

#### Supplementary Figure 8

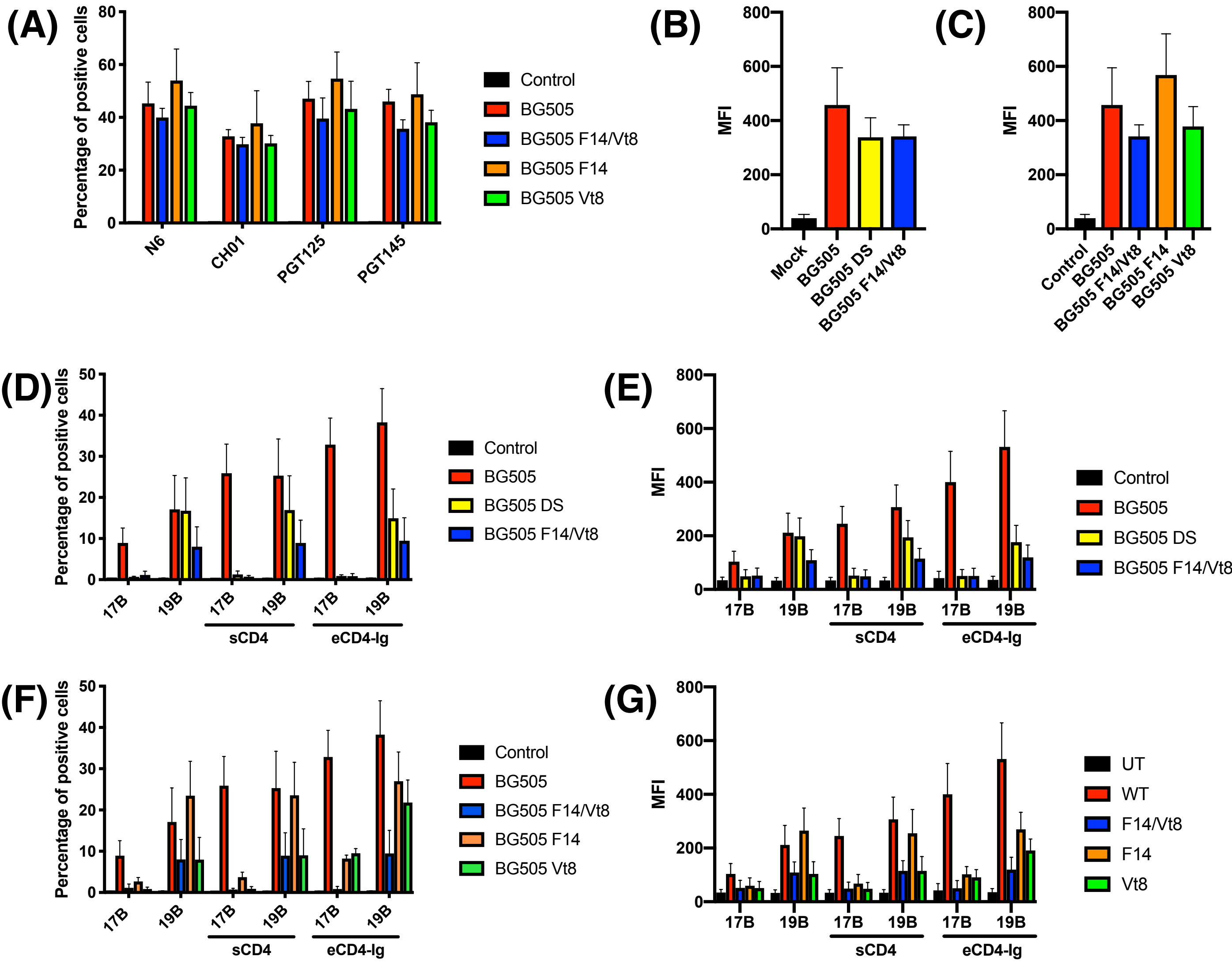

**Supplementary Figure 8**

**Supplementary Figure 8.** *Antigenicity and triggering of BG505 gp160 construct cell-surface expressed trimer.* ***(A)*** Percentage of cells positive for binding of bnAbs N6, CH01, PGT125, and PGT145 to 293F cell surface expressed gp160 BG505, BG505 F14/Vt8, BG505 F14, and BG505 Vt8 SOSIP trimers. ***(B)*** MFI data for binding of N6 to gp160 BG505, BG505 DS, and BG505 F14/Vt8 SOSIP trimers. ***(C)*** MFI data for binding of N6 to gp160 BG505, BG505 F14/Vt8, BG505 F14, and BG505 Vt8 SOSIP trimers. ***(D)*** Percentage of cells positive for binding of non-bnAbs 17B and 19B to 293F cell surface expressed BG505, BG505 DS, and BG505 F14/Vt8 SOSIP trimers in the presence and absence of sCD4 or CD4-Ig. ***(E)*** MFI data for binding of non-bnAbs 17B and 19B to 293F cell surface expressed BG505, BG505 DS, and BG505 F14/Vt8 SOSIP trimers in the presence and absence of sCD4 and CD4-Ig. ***(F)*** Percentage of cells positive for binding of non-bnAbs 17B and 19B to 293F cell surface expressed BG505, BG505 F14/Vt8, BG505 F14, and BG505 Vt8 SOSIP trimers in the presence and absence of sCD4 or CD4-Ig. ***(G)*** MFI data for binding of non-bnAbs 17B and 19B to 293F cell surface expressed BG505, BG505 F14/Vt8, BG505 F14, and BG505 Vt8 SOSIP trimers in the presence and absence of sCD4 and CD4-Ig.

**Supplementary Figure 9**

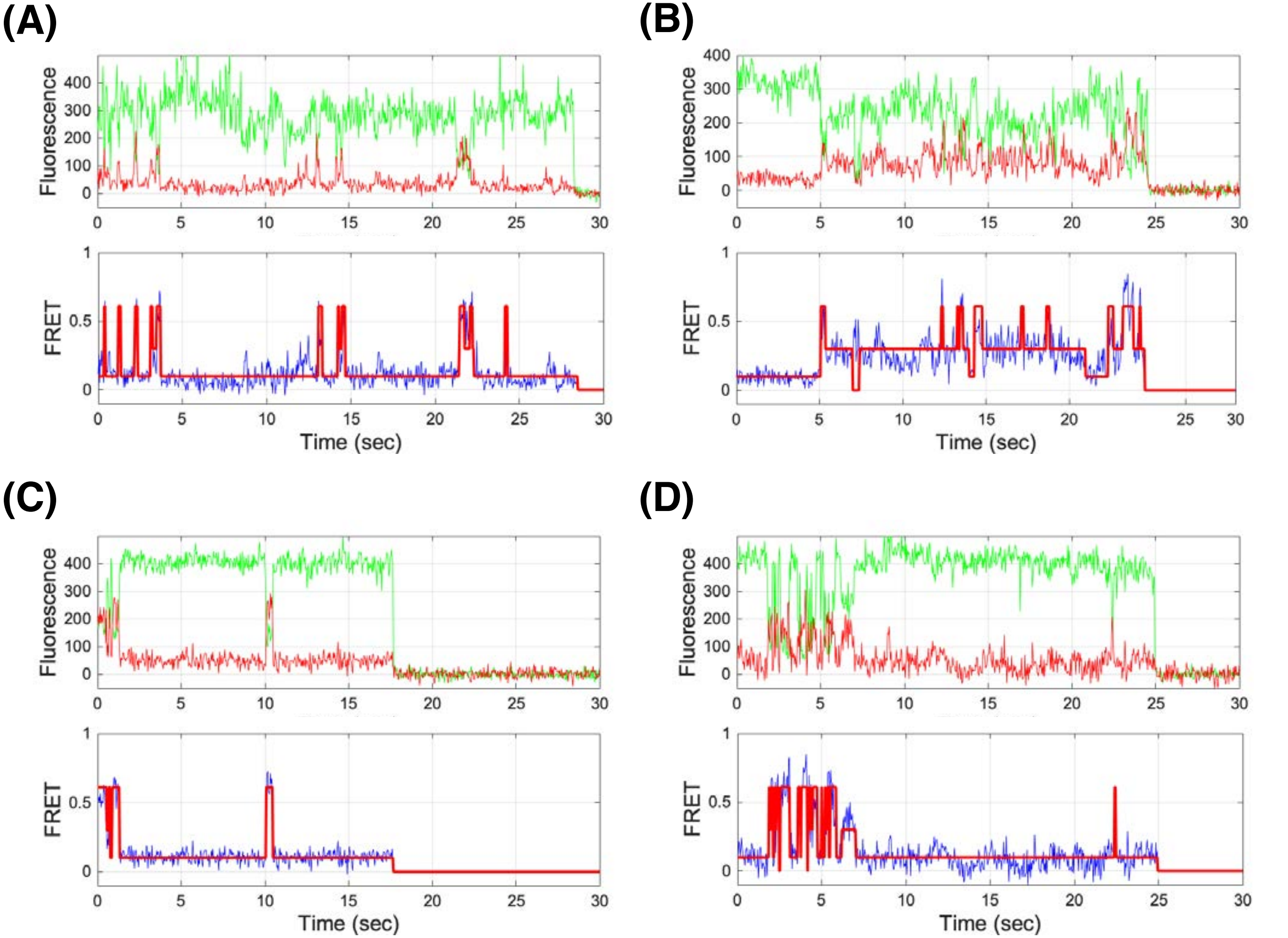


**Supplementary Figure 9.** *Representative fluorescence and FRET traces of both wild-type BG505 and BG505 F14/Vt8 virus Env in the absence and presence of 12xCD4* (Upper: Cy3 in green, Cy5 in red; Lower: resulting FRET in blue, HMM idealization in red) **(A)** Results for the BG505 Env. **(B)** Results for the BG505 Env in the presence of 12xCD4. **(C)** Results for the BG505 F14/Vt8 mutant Env. **(D)** Results for the BG505 F14/Vt8 Env mutant in the presence of 12xCD4.
